## Appendices for "Directional and disruptive selection in populations structured by class and continuous ontogeny under incomplete plasticity"

**Appendix A - Main results** **53**

### Appendix A - Main results

#### A.1 The basic reproductive number

We start by providing a representation of the basic reproductive number $R_0(\boldsymbol{u}, \boldsymbol{v})$ that is useful to do differential calculus, and do so by following previous calculations along the same lines (Lehmann et al. 2016; Ohtsuki et al. 2020). To simplify notation, we bring upfront that the reproductive number is a function of the mutant traits $\boldsymbol{z}(\boldsymbol{u}, \boldsymbol{v})$ and resident traits $\boldsymbol{v}$ (e.g., recall eqs. 1–2). Slightly abusing notation, in this appendix we write the reproductive number with phenotypic arguments as $R_0(\boldsymbol{z}, \boldsymbol{v})$ where it should be understood that $\boldsymbol{z} = \boldsymbol{z}(\boldsymbol{u}, \boldsymbol{v})$ and $\boldsymbol{v} = \boldsymbol{z}(\boldsymbol{v}, \boldsymbol{v})$.

We know from standard results (Mode 1969; Durinx et al. 2008; Metz & Leimar 2011; Inaba 2017) that $R_0(\boldsymbol{z}, \boldsymbol{v})$ is the leading eigenvalue of the next-generation matrix

$$\mathbf{R}(\boldsymbol{z}, \boldsymbol{v}) = \mathbf{R}(\boldsymbol{u}, \boldsymbol{v}) = \begin{bmatrix} \gamma_{11} R_{11}(\boldsymbol{u}_1, \boldsymbol{v}) & \dots & \gamma_{1n_c} R_{1n_c}(\boldsymbol{u}_{n_c}, \boldsymbol{v}) \\ \vdots & \ddots & \vdots \\ \gamma_{n_c1} R_{n_c1}(\boldsymbol{u}_1, \boldsymbol{v}) & \dots & \gamma_{n_cn_c} R_{n_cn_c}(\boldsymbol{u}_{n_c}, \boldsymbol{v}) \end{bmatrix}, \tag{A-1}$$

satisfying

$$R_0(\boldsymbol{z}, \boldsymbol{v}) \boldsymbol{q}(\boldsymbol{z}, \boldsymbol{v}) = \mathbf{R}(\boldsymbol{z}, \boldsymbol{v}) \boldsymbol{q}(\boldsymbol{z}, \boldsymbol{v}), \tag{A-2}$$

where $\boldsymbol{q}(\boldsymbol{z}, \boldsymbol{v})$ is the right-eigenvector of $\mathbf{R}(\boldsymbol{z}, \boldsymbol{v})$, that we always scale so that its elements sum to 1. In a monomorphic population at equilibrium, we know that $R_0(\boldsymbol{v}, \boldsymbol{v}) = 1$. Thus, the leading left-eigenvector $\boldsymbol{\nu}^\circ$ of $\mathbf{R}^\circ = \mathbf{R}(\boldsymbol{v}, \boldsymbol{v})$ is the row vector satisfying

$$\boldsymbol{\nu}^\circ = \boldsymbol{\nu}^\circ \mathbf{R}^\circ. \tag{A-3}$$

Thorough, we scale $\boldsymbol{\nu}^\circ$ so that $\boldsymbol{\nu}^\circ \cdot \boldsymbol{q}^\circ = 1$, where the dot notation refers to the standard dot product, which we use exclusively to indicate the inner product of two vectors (i.e. yielding a scalar). Left-multiplying both sides of eq. (A-2) by $\boldsymbol{\nu}^\circ$ and re-arranging gives

$$R_0(\boldsymbol{z}, \boldsymbol{v}) = \frac{\boldsymbol{\nu}^\circ \mathbf{R}(\boldsymbol{z}, \boldsymbol{v}) \boldsymbol{q}(\boldsymbol{z}, \boldsymbol{v})}{\boldsymbol{\nu}^\circ \cdot \boldsymbol{q}(\boldsymbol{z}, \boldsymbol{v})}. \tag{A-4}$$

Let us denote by

$$\delta_{ij}(\boldsymbol{z}_j, \boldsymbol{v}) = R_{ij}(\boldsymbol{z}_j, \boldsymbol{v}) - R_{ij}(\boldsymbol{v}_j, \boldsymbol{v}) = R_{ij}(\boldsymbol{z}_j, \boldsymbol{v}) - R_{ij}^\circ. \tag{A-5}$$

the difference in the expected lifetime numbers of offspring of class $i$ produced by mutants and residents of class $j$ (given by eq. 1). Expressing eq. (A-4) in terms of this difference gives

$$
\begin{aligned}
R_0(\boldsymbol{z},\boldsymbol{v}) &= \frac{1}{\boldsymbol{\nu}^\circ\cdot\boldsymbol{q}(\boldsymbol{z},\boldsymbol{v})}\left[\sum_{i=1}^{n_\mathrm{c}}\sum_{j=1}^{n_\mathrm{c}}\nu_i^\circ\gamma_{ij}\left(R_{ij}^\circ+\delta_{ij}(\boldsymbol{z}_j,\boldsymbol{v})\right)q_j(\boldsymbol{z},\boldsymbol{v})\right] \\
&= \frac{1}{\boldsymbol{\nu}^\circ\cdot\boldsymbol{q}(\boldsymbol{z},\boldsymbol{v})}\left[\sum_{j=1}^{n_\mathrm{c}}\sum_{i=1}^{n_\mathrm{c}}\nu_i^\circ\gamma_{ij}R_{ij}^\circ q_j(\boldsymbol{z},\boldsymbol{v})\right] \\
&\quad+\frac{1}{\boldsymbol{\nu}^\circ\cdot\boldsymbol{q}(\boldsymbol{z},\boldsymbol{v})}\left[\sum_{i=1}^{n_\mathrm{c}}\sum_{j=1}^{n_\mathrm{c}}\nu_i^\circ\gamma_{ij}\delta_{ij}(\boldsymbol{z}_j,\boldsymbol{v})q_j(\boldsymbol{z},\boldsymbol{v})\right].
\end{aligned}
\tag{A-6}
$$

Since $\sum_{i=1}^{n_\mathrm{c}}\nu_i^\circ\gamma_{ij}R_{ij}^\circ = \nu_j^\circ$ (by virtue of eq. A-3) and since $\sum_{j=1}^{n_\mathrm{c}}\nu_j^\circ q_j(\boldsymbol{z},\boldsymbol{v}) = \boldsymbol{\nu}^\circ\cdot\boldsymbol{q}(\boldsymbol{z},\boldsymbol{v})$, eq. (A-6) simplifies to

$$
R_0(\boldsymbol{z},\boldsymbol{v}) = 1+\frac{1}{\boldsymbol{\nu}^\circ\cdot\boldsymbol{q}(\boldsymbol{z},\boldsymbol{v})}\left[\sum_{i=1}^{n_\mathrm{c}}\sum_{j=1}^{n_\mathrm{c}}\nu_i^\circ\gamma_{ij}\delta_{ij}(\boldsymbol{z}_j,\boldsymbol{v})q_j(\boldsymbol{z},\boldsymbol{v})\right]. \tag{A-7}
$$

We will use eq. (A-7) as a starting point to obtain expressions for derivatives of the basic reproductive number.

#### A.2 Directional selection

##### A.2.1 Lifetime selection gradients

Taking the derivative of eq. (A-7) with respect to $z_{j,l}$, applying the product rule of differentiation twice and evaluating at the resident allelic values gives

$$
\begin{aligned}
\left.\frac{\partial R_0(\boldsymbol{z},\boldsymbol{v})}{\partial z_{j,l}}\right|_{\boldsymbol{z}=\boldsymbol{v}} &= \frac{1}{\boldsymbol{\nu}^\circ\cdot\boldsymbol{q}(\boldsymbol{v},\boldsymbol{v})}\left[\sum_{m=1}^{n_\mathrm{c}}\sum_{n=1}^{n_\mathrm{c}}\nu_m^\circ\gamma_{mn}\left.\frac{\partial\delta_{mn}(\boldsymbol{z}_n,\boldsymbol{v})}{\partial z_{j,l}}\right|_{\boldsymbol{z}=\boldsymbol{v}}q_n(\boldsymbol{v},\boldsymbol{v})\right] \\
&\quad+\frac{1}{\boldsymbol{\nu}^\circ\cdot\boldsymbol{q}(\boldsymbol{v},\boldsymbol{v})}\left[\sum_{m=1}^{n_\mathrm{c}}\sum_{n=1}^{n_\mathrm{c}}\nu_m^\circ\gamma_{mn}\underbrace{\delta_{mn}^\circ}_{0}\left.\frac{\partial q_n(\boldsymbol{z},\boldsymbol{v})}{\partial z_{j,l}}\right|_{\boldsymbol{z}=\boldsymbol{v}}\right] \\
&\quad+\left.\frac{\partial(\boldsymbol{\nu}^\circ\cdot\boldsymbol{q}(\boldsymbol{z},\boldsymbol{v}))^{-1}}{\partial z_{j,l}}\right|_{\boldsymbol{z}=\boldsymbol{v}}\left[\sum_{m=1}^{n_\mathrm{c}}\sum_{n=1}^{n_\mathrm{c}}\nu_m^\circ\gamma_{mn}\underbrace{\delta_{mn}^\circ}_{0}q_n(\boldsymbol{v},\boldsymbol{v})\right] \\
&= \frac{1}{\boldsymbol{\nu}^\circ\cdot\boldsymbol{q}^\circ}\left[\sum_{m=1}^{n_\mathrm{c}}\sum_{n=1}^{n_\mathrm{c}}\nu_m^\circ\gamma_{mn}\left.\frac{\partial\delta_{mn}(\boldsymbol{z}_n,\boldsymbol{v})}{\partial z_{j,l}}\right|_{\boldsymbol{z}=\boldsymbol{v}}q_n^\circ\right].
\end{aligned}
\tag{A-8}
$$

Recalling that $\boldsymbol{\nu}^\circ\cdot\boldsymbol{q}^\circ = 1$, replacing derivatives of $\delta_{mn}$ by derivatives of $R_{mn}$ (as these are equal according to eq. A-5) and plugging back into (3b) shows that the selection gradient specific to the $l$th trait expressed in class $j$ is

$$\begin{aligned} s_{j,l}(\boldsymbol{v}) &= \left[\sum_{m=1}^{n_c}\sum_{n=1}^{n_c} \nu_m^\circ \gamma_{mn} \left.\frac{\partial R_{mn}(\boldsymbol{z}_n, \boldsymbol{v})}{\partial z_{j,l}}\right|_{\boldsymbol{z}=\boldsymbol{v}} q_n^\circ\right] \left.\frac{\partial z_{j,l}(u_{j,l}, v_{j,l})}{\partial u_{j,l}}^\circ\right|_{\boldsymbol{u}=\boldsymbol{v}} \\ &= \left[\sum_{m=1}^{n_c} \nu_m^\circ \gamma_{mj} \left.\frac{\partial R_{mj}(\boldsymbol{z}_j, \boldsymbol{v})}{\partial z_{j,l}}\right|_{\boldsymbol{z}=\boldsymbol{v}} q_j^\circ\right] \left.\frac{\partial z_{j,l}(u_{j,l}, v_{j,l})}{\partial u_{j,l}}\right|_{\boldsymbol{u}=\boldsymbol{v}}, \end{aligned} \tag{A-9}$$

where one sum was removed since $\partial R_{mn}(\boldsymbol{z}_n, \boldsymbol{v})/\partial z_{j,l} = 0$ for all $n \neq j$. Changing variable $m$ to $i$, we finally have

$$s_{j,l}(\boldsymbol{v}) = \left[\sum_{i=1}^{n_c} \nu_i^\circ \gamma_{ij} \left.\frac{\partial R_{ij}(\boldsymbol{z}_j, \boldsymbol{v})}{\partial z_{j,l}}\right|_{\boldsymbol{z}=\boldsymbol{v}} q_j^\circ\right] \left.\frac{\partial z_{j,l}(u_{j,l}, v_{j,l})}{\partial u_{j,l}}\right|_{\boldsymbol{u}=\boldsymbol{v}}. \tag{A-10}$$

##### A.2.2 Hamiltonians and age-specific directional selection

We now derive eq. (8) of the main text, drawing inspiration from calculations done in optimal control theory (e.g., Liberzon 2012; Kamien & Schwartz 2013) as applied to constant controls (Lehmann 2022), which we here generalize to class-structured populations. To that end, it is useful to treat the class-specific survival to age $a$, $l_j(a)$, in the same way as the internal states of individuals of class $j$. We thus define $\boldsymbol{x}_j(a) = (l_j(a), \boldsymbol{x}_j(a))$ as the vector of size $(n_x + 1)$ concatenating the survival probability and the internal states in class $j$, and $\tilde{\boldsymbol{g}}_j(\boldsymbol{z}_j, \boldsymbol{x}_j(a), \boldsymbol{v}) = (-\mu_j(\boldsymbol{z}_j, \boldsymbol{x}_j(a), \boldsymbol{v}) l_j(a), \boldsymbol{g}_j(\boldsymbol{z}_j, \boldsymbol{x}_j(a), \boldsymbol{v}))$ as the vector of size $(n_x + 1)$ concatenating the rates of change in survivorship and internal states in class $j$. For brevity, we refer to all elements of $\boldsymbol{x}_j(a)$ as *states* and use the term *internal states* to refer specifically to the variables in $\boldsymbol{x}_j(a)$ when needed.

With these definitions, we start by adding a zero term to the integrand of eq. (2), and write

$$R_{ij}(\boldsymbol{z}_j, \boldsymbol{v}) = \int_0^\infty \left[l_j(a) f_{ij}(\boldsymbol{z}_j, \boldsymbol{x}_j(a), \boldsymbol{v}) + \tilde{\boldsymbol{\lambda}}_{ij}(a) \cdot \underbrace{\left(\tilde{\boldsymbol{g}}_j(\boldsymbol{z}_j, \boldsymbol{x}_j(a), \boldsymbol{v}) - \frac{\mathrm{d}\boldsymbol{x}_j(a)}{\mathrm{d}a}\right)}_{0}\right] \mathrm{d}a, \tag{A-11}$$

where $\tilde{\boldsymbol{\lambda}}_{ij}(a) = (\lambda_{ij}^{\mathrm{l}}(a), \boldsymbol{\lambda}_{ij}^{\mathrm{x}}(a))$ is a vector of costates. Therein, $\lambda_{ij}^{\mathrm{l}}(a)$ is the costate associated with survivorship, and $\boldsymbol{\lambda}_{ij}^{\mathrm{x}}(a) = (\lambda_{ij,1}^{\mathrm{x}}, ..., \lambda_{ij,n_x}^{\mathrm{x}})$ is a vector of size $n_x$ collecting the costates associated with internal states. Using the class-oriented Hamiltonian, given by

$$H_{ij}(\boldsymbol{z}_j, \boldsymbol{x}_j(a), \boldsymbol{v}) = l_j(a) f_{ij}(\boldsymbol{z}_j, \boldsymbol{x}_j(a), \boldsymbol{v}) + \tilde{\boldsymbol{\lambda}}_{ij}(a) \cdot \tilde{\boldsymbol{g}}_j(\boldsymbol{z}_j, \boldsymbol{x}_j(a), \boldsymbol{v}), \tag{A-12}$$

eq. (A-11) becomes

$$R_{ij}(\boldsymbol{z}_j, \boldsymbol{v}) = \int_0^\infty \left[H_{ij}(\boldsymbol{z}_j, \boldsymbol{x}_j(a), \boldsymbol{v}) - \tilde{\boldsymbol{\lambda}}_{ij}(a) \cdot \frac{\mathrm{d}\boldsymbol{x}_j(a)}{\mathrm{d}a}\right] \mathrm{d}a. \tag{A-13}$$

The last term in the integrand of eq. (A-13) can then be integrated by parts to give

$$\int_0^\infty \tilde{\boldsymbol{\lambda}}_{ij}(a) \cdot \frac{\mathrm{d}\boldsymbol{x}_j(a)}{\mathrm{d}a} \mathrm{d}a = \tilde{\boldsymbol{\lambda}}_{ij}(\infty) \cdot \boldsymbol{x}_j(\infty) - \tilde{\boldsymbol{\lambda}}_{ij}(0) \cdot \boldsymbol{x}_j(0) - \int_0^\infty \frac{\mathrm{d}\tilde{\boldsymbol{\lambda}}_{ij}(a)}{\mathrm{d}a} \cdot \boldsymbol{x}_j(a) \mathrm{d}a. \tag{A-14}$$

Thus, we have

$$\begin{aligned} R_{ij}(\boldsymbol{z}_j, \boldsymbol{v}) = \int_0^\infty \left[ H_{ij}(\boldsymbol{z}_j, \boldsymbol{x}_j(a), \boldsymbol{v}) + \frac{\mathrm{d}\tilde{\boldsymbol{\lambda}}_{ij}(a)}{\mathrm{d}a} \cdot \boldsymbol{x}_j(a) \right] \mathrm{d}a \\ - \tilde{\boldsymbol{\lambda}}_{ij}(\infty) \cdot \boldsymbol{x}_j(\infty) + \tilde{\boldsymbol{\lambda}}_{ij}(0) \cdot \boldsymbol{x}_j(0). \end{aligned} \tag{A-15}$$

We can now take the derivative of eq. (A-15) with respect to $z_{j,l}$. Making use of the Leibniz formula and the chain rule, re-labeling dummy variables and recalling that mutant state values $\boldsymbol{x}_n(a)$ depend on mutant trait values, we obtain

$$\begin{aligned} \left.\frac{\partial R_{mn}(\boldsymbol{z}_n, \boldsymbol{v})}{\partial z_{j,l}}\right|_{\boldsymbol{z}=\boldsymbol{v}} = \int_0^\infty & \left[ \left.\frac{\partial H_{mn}(\boldsymbol{z}_n, \boldsymbol{x}_n^\circ(a), \boldsymbol{v})}{\partial z_{j,l}}\right|_{\boldsymbol{z}=\boldsymbol{v}} \right. \\ & \left. + \left( \left.\frac{\partial H_{mn}(\boldsymbol{z}_n, \boldsymbol{x}_n(a), \boldsymbol{v})}{\partial \boldsymbol{x}_n(a)}\right|_{\boldsymbol{z}=\boldsymbol{v}} + \frac{\mathrm{d}\tilde{\boldsymbol{\lambda}}_{mn}(a)}{\mathrm{d}a} \right) \cdot \left.\frac{\partial \boldsymbol{x}_n(a)}{\partial z_{j,l}}\right|_{\boldsymbol{z}=\boldsymbol{v}} \right] \mathrm{d}a \\ & - \tilde{\boldsymbol{\lambda}}_{mn}(\infty) \cdot \left.\frac{\partial \boldsymbol{x}_n(\infty)}{\partial z_{j,l}}\right|_{\boldsymbol{z}=\boldsymbol{v}}, \end{aligned} \tag{A-16}$$

where initial state values $\boldsymbol{x}_n(0)$ were, by assumption, independent of trait values, such that their derivatives with respect to $z_{j,l}$ vanished. Note that we slightly abused notation in the first line of eq. (A-16), pre-emptively substituting state values $\boldsymbol{x}_n(a)$ with resident state values $\boldsymbol{x}_n^\circ(a)$, to make more explicit the fact that the derivative of the Hamiltonian with respect to the mutant trait is a direct derivative (i.e. the indirect effects of the trait via internal state values are accounted for by costates in the second line of the equation).

We can see from eq. (A-16) that if costates are chosen such that

$$\frac{\mathrm{d}\tilde{\boldsymbol{\lambda}}_{mn}(a)}{\mathrm{d}a} = - \left.\frac{\partial H_{mn}(\boldsymbol{z}_n, \boldsymbol{x}_n(a), \boldsymbol{v})}{\partial \boldsymbol{x}_n(a)}\right|_{\boldsymbol{z}=\boldsymbol{v}} \quad \text{and} \quad \tilde{\boldsymbol{\lambda}}_{mn}(\infty) \cdot \left.\frac{\partial \boldsymbol{x}_n(\infty)}{\partial z_{j,l}}\right|_{\boldsymbol{z}=\boldsymbol{v}} = 0 \tag{A-17}$$

then eq. (A-18) simplifies to

$$\left.\frac{\partial R_{mn}(\boldsymbol{z}_n, \boldsymbol{v})}{\partial z_{j,l}}\right|_{\boldsymbol{z}=\boldsymbol{v}} = \int_0^\infty \left.\frac{\partial H_{mn}(\boldsymbol{z}_n, \boldsymbol{x}_n(a), \boldsymbol{v})}{\partial z_{j,l}}\right|_{\boldsymbol{z}=\boldsymbol{v}} \mathrm{d}a. \tag{A-18}$$

##### A.2.3 Boundary conditions for costates

In order to completely characterize the dynamical systems that determine costates, we now use eq. (A-17) to obtain boundary conditions for these functions.

It is straightforward to obtain boundary conditions for the costates associated to internal states, using the second condition in eq. (A-17). We know that, in general, it holds that $\partial \boldsymbol{x}_n(\infty)/\partial z_{j,l}|_{\boldsymbol{z}=\boldsymbol{v}} > 0$ since traits can affect development at any point of life history. Thus, for eq. (A-17) to be true, it must be that all costates associated to internal states converge to 0,

$$\lim_{a\to\infty} \boldsymbol{\lambda}^{\mathrm{x}}_{mn}(a) = \mathbf{0}. \tag{A-19}$$

Finding boundary conditions for costates associated to survivorship is less straightforward. This is because we have $\partial l_n(\infty)/\partial z_{j,l}|_{\boldsymbol{z}=\boldsymbol{v}} = 0$, as $\lim_{a\to\infty} l_n(a) = 0$ (see also eq. A-49 with $a \to \infty$), meaning that eq. (A-17) does not impose the condition that $\lambda^{\mathrm{l}}_{mn}(\infty) = 0$. To obtain a boundary condition, we can rely instead on the dynamical equation

$$\frac{\mathrm{d}\lambda^{\mathrm{l}}_{mn}(a)}{\mathrm{d}a} = -f_{mn}(\boldsymbol{v}_n, \boldsymbol{x}^{\circ}_n(a), \boldsymbol{v}) + \lambda^{\mathrm{l}}_{mn}(a)\mu_n(\boldsymbol{v}_n, \boldsymbol{x}^{\circ}_n(a), \boldsymbol{v}), \tag{A-20}$$

obtained from eq. (A-17). Using the generalized Cauchy formula (Weber 2011; see eq. A-47 in appendix A.4.1) with $\zeta = 0$, eq. (A-20) implies that $\lambda^{\mathrm{l}}_{mn}(a)$ can be written

$$\lambda^{\mathrm{l}}_{mn}(a) = \tilde{\Psi}_n(a, 0)\lambda^{\mathrm{l}}_{mn}(0) - \int_0^a \tilde{\Psi}_n(a, \tau) f_{mn}(\boldsymbol{v}_n, \boldsymbol{x}^{\circ}_n(\tau), \boldsymbol{v})\mathrm{d}\tau, \tag{A-21}$$

where $\tilde{\Psi}_n(a, \tau)$ is the solution to

$$\frac{\mathrm{d}\tilde{\Psi}_n(a, \tau)}{\mathrm{d}a} = \mu_n(\boldsymbol{v}_n, \boldsymbol{x}^{\circ}_n(a), \boldsymbol{v})\tilde{\Psi}_n(a, \tau) \text{ with i.c. } \tilde{\Psi}_n(\tau, \tau) = 1, \tag{A-22}$$

following eq. (A-48). Eq. (A-22) admits the following solution,

$$\begin{aligned}
\tilde{\Psi}_n(a, \tau) &= \exp\left[\int_\tau^a \mu_n(\boldsymbol{v}_n, \boldsymbol{x}^{\circ}_n(s), \boldsymbol{v})\mathrm{d}s\right] \\
&= \exp\left[-\int_a^\tau \mu_n(\boldsymbol{v}_n, \boldsymbol{x}^{\circ}_n(s), \boldsymbol{v})\mathrm{d}s\right] \\
&= \exp\left[-\int_0^\tau \mu_n(\boldsymbol{v}_n, \boldsymbol{x}^{\circ}_n(s), \boldsymbol{v})\mathrm{d}s\right] \Big/ \exp\left[-\int_0^a \mu_n(\boldsymbol{v}_n, \boldsymbol{x}^{\circ}_n(s), \boldsymbol{v})\mathrm{d}s\right] \\
&= \frac{l^{\circ}_n(\tau)}{l^{\circ}_n(a)},
\end{aligned} \tag{A-23}$$

where we used the fact that $l^{\circ}_n(a) = \exp\left[-\int_0^a \mu_n(\boldsymbol{v}_n, \boldsymbol{x}^{\circ}_n(s), \boldsymbol{v})\mathrm{d}s\right]$ is the unique solution to the first line of eq. (12). Plugging eq. (A-23) into eq. (A-21) and with $l^{\circ}_n(0) = 1$, rearranging and using eq. (1) gives

$$\begin{aligned}
\lambda^{\mathrm{l}}_{mn}(a) &= \frac{1}{l^{\circ}_n(a)}\left[\lambda^{\mathrm{l}}_{mn}(0) - \int_0^a l^{\circ}_n(\tau) f_{mn}(\boldsymbol{v}_n, \boldsymbol{x}^{\circ}_n(\tau), \boldsymbol{v})\mathrm{d}\tau\right] \\
&= \frac{1}{l^{\circ}_n(a)}\left[\lambda^{\mathrm{l}}_{mn}(0) - \int_0^\infty l^{\circ}_n(\tau) f_{mn}(\boldsymbol{v}_n, \boldsymbol{x}^{\circ}_n(\tau), \boldsymbol{v})\mathrm{d}\tau + \int_a^\infty l^{\circ}_n(\tau) f_{mn}(\boldsymbol{v}_n, \boldsymbol{x}^{\circ}_n(\tau), \boldsymbol{v})\mathrm{d}\tau\right] \\
&= \frac{1}{l^{\circ}_n(a)}\left[\lambda^{\mathrm{l}}_{mn}(0) - R^{\circ}_{ij}\right] + \frac{1}{l^{\circ}_n(a)}\int_a^\infty l^{\circ}_n(\tau) f_{mn}(\boldsymbol{v}_n, \boldsymbol{x}^{\circ}_n(\tau), \boldsymbol{v})\mathrm{d}\tau.
\end{aligned} \tag{A-24}$$

Since $\lim_{a\to\infty} l^{\circ}_n(a) = 0$, the first term in the last line of eq. (A-24) will tend to infinity, as $a \to \infty$, if $\lambda^{\mathrm{l}}_{mn}(0) - R^{\circ}_{ij}$ is non-zero. As reproduction cannot be infinite, biological consistency thus entails the initial condition

$$\lambda^{\mathrm{l}}_{mn}(0) = R^{\circ}_{ij}. \tag{A-25}$$

Note that with such initial condition, eq. (A-24) becomes eq. (I-C),

$$\lambda^{\mathrm{l}}_{mn}(a) = \frac{1}{l^{\circ}_{n}(a)} \int_{a}^{\infty} l^{\circ}_{n}(\tau) f_{mn}(\boldsymbol{v}_n, \boldsymbol{x}^{\circ}_{n}(\tau), \boldsymbol{v})\mathrm{d}\tau, \tag{A-26}$$

showing that $\lambda^{\mathrm{l}}_{mn}(a)$ is equivalent to Fisher's reproductive value in class-structured populations (see box 1).

#### A.3 Quadratic selection

In this section we derive eqs. (14)–(20) of the main text.

##### A.3.1 Lifetime Hessian matrices

We start by taking the derivative of eq. (A-7) with respect to $z_{i,k}$ and $z_{j,l}$. Applying the product rule and evaluating at resident trait values, we have

$$\begin{aligned}
\left.\frac{\partial^2 R_0(\boldsymbol{z},\boldsymbol{v})}{\partial z_{i,k}\partial z_{j,l}}\right|_{\boldsymbol{z}=\boldsymbol{v}} &= \frac{1}{\boldsymbol{\nu}^{\circ}\cdot\boldsymbol{q}^{\circ}} \left[\sum_{m=1}^{n_{\mathrm{c}}}\sum_{n=1}^{n_{\mathrm{c}}} \nu^{\circ}_{m}\gamma_{mn} \left.\left(\frac{\partial \delta_{mn}(\boldsymbol{z}_n,\boldsymbol{v})}{\partial z_{i,k}} \frac{\partial q_n(\boldsymbol{z},\boldsymbol{v})}{\partial z_{j,l}}\right)\right|_{\boldsymbol{z}=\boldsymbol{v}}\right] \\
&+ \frac{1}{\boldsymbol{\nu}^{\circ}\cdot\boldsymbol{q}^{\circ}} \left[\sum_{m=1}^{n_{\mathrm{c}}}\sum_{n=1}^{n_{\mathrm{c}}} \nu^{\circ}_{m}\gamma_{mn} \left.\left(\frac{\partial \delta_{mn}(\boldsymbol{z}_n,\boldsymbol{v})}{\partial z_{j,l}} \frac{\partial q_n(\boldsymbol{z},\boldsymbol{v})}{\partial z_{i,k}}\right)\right|_{\boldsymbol{z}=\boldsymbol{v}}\right] \\
&+ \frac{1}{\boldsymbol{\nu}^{\circ}\cdot\boldsymbol{q}^{\circ}} \left[\sum_{m=1}^{n_{\mathrm{c}}}\sum_{n=1}^{n_{\mathrm{c}}} \nu^{\circ}_{m}\gamma_{mn} \left.\frac{\partial^2 \delta_{mn}(\boldsymbol{z}_n,\boldsymbol{v})}{\partial z_{i,k}\partial z_{j,l}}\right|_{\boldsymbol{z}=\boldsymbol{v}} q^{\circ}_{n}\right] \\
&+ \frac{1}{\boldsymbol{\nu}^{\circ}\cdot\boldsymbol{q}^{\circ}} \left[\sum_{m=1}^{n_{\mathrm{c}}}\sum_{n=1}^{n_{\mathrm{c}}} \nu^{\circ}_{m}\gamma_{mn} \underbrace{\delta^{\circ}_{mn}}_{0} \left.\frac{\partial^2 q_n(\boldsymbol{z},\boldsymbol{v})}{\partial z_{i,k}\partial z_{j,l}}\right|_{\boldsymbol{z}=\boldsymbol{v}}\right] \\
&+ \left.\frac{\partial(\boldsymbol{\nu}^{\circ}\cdot\boldsymbol{q}(\boldsymbol{z},\boldsymbol{v}))^{-1}}{\partial z_{i,k}}\right|_{\boldsymbol{z}=\boldsymbol{v}} \left[\sum_{m=1}^{n_{\mathrm{c}}}\sum_{n=1}^{n_{\mathrm{c}}} \nu^{\circ}_{m}\gamma_{mn} \left.\frac{\partial \delta_{mn}(\boldsymbol{z}_n,\boldsymbol{v})}{\partial z_{j,l}}\right|_{\boldsymbol{z}=\boldsymbol{v}} q^{\circ}_{n}\right] \\
&+ \left.\frac{\partial(\boldsymbol{\nu}^{\circ}\cdot\boldsymbol{q}(\boldsymbol{z},\boldsymbol{v}))^{-1}}{\partial z_{j,l}}\right|_{\boldsymbol{z}=\boldsymbol{v}} \left[\sum_{m=1}^{n_{\mathrm{c}}}\sum_{n=1}^{n_{\mathrm{c}}} \nu^{\circ}_{m}\gamma_{mn} \left.\frac{\partial \delta_{mn}(\boldsymbol{z}_n,\boldsymbol{v})}{\partial z_{i,k}}\right|_{\boldsymbol{z}=\boldsymbol{v}} q^{\circ}_{n}\right] \\
&+ \left.\frac{\partial(\boldsymbol{\nu}^{\circ}\cdot\boldsymbol{q}(\boldsymbol{z},\boldsymbol{v}))^{-1}}{\partial z_{i,k}}\right|_{\boldsymbol{z}=\boldsymbol{v}} \left[\sum_{m=1}^{n_{\mathrm{c}}}\sum_{n=1}^{n_{\mathrm{c}}} \nu^{\circ}_{m}\gamma_{mn} \underbrace{\delta^{\circ}_{mn}}_{0} \left.\frac{\partial q_n(\boldsymbol{z},\boldsymbol{v})}{\partial z_{j,l}}\right|_{\boldsymbol{z}=\boldsymbol{v}}\right] \\
&+ \left.\frac{\partial(\boldsymbol{\nu}^{\circ}\cdot\boldsymbol{q}(\boldsymbol{z},\boldsymbol{v}))^{-1}}{\partial z_{j,l}}\right|_{\boldsymbol{z}=\boldsymbol{v}} \left[\sum_{m=1}^{n_{\mathrm{c}}}\sum_{n=1}^{n_{\mathrm{c}}} \nu^{\circ}_{m}\gamma_{mn} \underbrace{\delta^{\circ}_{mn}}_{0} \left.\frac{\partial q_n(\boldsymbol{z},\boldsymbol{v})}{\partial z_{i,k}}\right|_{\boldsymbol{z}=\boldsymbol{v}}\right] \\
&+ \left.\frac{\partial^2(\boldsymbol{\nu}^{\circ}\cdot\boldsymbol{q}(\boldsymbol{z},\boldsymbol{v}))^{-1}}{\partial z_{i,k}\partial z_{j,l}}\right|_{\boldsymbol{z}=\boldsymbol{v}} \left[\sum_{m=1}^{n_{\mathrm{c}}}\sum_{n=1}^{n_{\mathrm{c}}} \nu^{\circ}_{m}\gamma_{mn} \underbrace{\delta^{\circ}_{mn}}_{0} q^{\circ}_{n}\right].
\end{aligned} \tag{A-27}$$

The first two lines of eq. (A-27) can be simplified by removing a sum, since $\partial\delta_{mn}(\boldsymbol{z}_n,\boldsymbol{v})/\partial z_{j,l}=0$ for all $j\neq n$. In addition, we know that $\partial(\boldsymbol{\nu}^\circ\cdot\boldsymbol{q}(\boldsymbol{z},\boldsymbol{v}))^{-1}/\partial z_{j,l}|_{\boldsymbol{z}=\boldsymbol{v}} = -\boldsymbol{\nu}^\circ\cdot(\partial\boldsymbol{q}(\boldsymbol{z},\boldsymbol{v})/\partial z_{j,l}|_{\boldsymbol{z}=\boldsymbol{v}})$ since $(\boldsymbol{\nu}^\circ\cdot\boldsymbol{q}^\circ)^{-2}=1$ owing to the fact that $\boldsymbol{\nu}^\circ\cdot\boldsymbol{q}^\circ=1$. With these simplifications, replacing derivatives of $\delta_{mn}$ by derivatives of $R_{mn}$ (as these are equal by definition; see eq. A-5), using eqs. (3b) and (A-9), and plugging eq. (A-27) back into eq. (3c) gives element $(k,l)$ of the class-oriented Hessian matrix $\mathbf{H}_{ij}(\boldsymbol{v})$ as

$$\begin{aligned} h_{ij,kl}(\boldsymbol{v}) = &\left[\sum_{m=1}^{n_c}\sum_{n=1}^{n_c}\nu_m^\circ\gamma_{mn}\left.\frac{\partial^2 R_{mn}(\boldsymbol{z}_n,\boldsymbol{v})}{\partial z_{i,k}\partial z_{j,l}}\right|_{\boldsymbol{z}=\boldsymbol{v}} q_n^\circ\right]\times\left.\frac{\partial z_{i,k}(u_{i,k},v_{i,k})}{\partial u_{i,k}}\right|_{\boldsymbol{u}=\boldsymbol{v}}\times\left.\frac{\partial z_{j,l}(u_{j,l},v_{j,l})}{\partial u_{j,l}}\right|_{\boldsymbol{u}=\boldsymbol{v}} \\ &+h_{ij,kl}^{\mathrm{zq}}(\boldsymbol{v})-h_{ij,kl}^{\mathrm{q}}(\boldsymbol{v}), \end{aligned} \tag{A-28}$$

where

$$\begin{aligned} h_{ij,kl}^{\mathrm{zq}}(\boldsymbol{v}) = &\left[\sum_{m=1}^{n_c}\nu_m^\circ\left.\left(\gamma_{mi}\frac{\partial R_{mi}(\boldsymbol{z}_i,\boldsymbol{v})}{\partial z_{i,k}}\frac{\partial q_i(\boldsymbol{z},\boldsymbol{v})}{\partial z_{j,l}}+\gamma_{mj}\frac{\partial R_{mj}(\boldsymbol{z}_j,\boldsymbol{v})}{\partial z_{j,l}}\frac{\partial q_j(\boldsymbol{z},\boldsymbol{v})}{\partial z_{i,k}}\right)\right|_{\boldsymbol{z}=\boldsymbol{v}}\right] \\ &\times\left.\frac{\partial z_{i,k}(u_{i,k},v_{i,k})}{\partial u_{i,k}}\right|_{\boldsymbol{u}=\boldsymbol{v}}\times\left.\frac{\partial z_{j,l}(u_{j,l},v_{j,l})}{\partial u_{j,l}}\right|_{\boldsymbol{u}=\boldsymbol{v}}, \end{aligned} \tag{A-29}$$

and where the term

$$\begin{aligned} h_{ij,kl}^{\mathrm{q}}(\boldsymbol{v}) = &\left(\boldsymbol{\nu}^\circ\cdot\left.\frac{\partial\boldsymbol{q}(\boldsymbol{z},\boldsymbol{v})}{\partial z_{i,k}}\right|_{\boldsymbol{z}=\boldsymbol{v}}\right)\times s_{j,l}(\boldsymbol{v})\times\left.\frac{\partial z_{i,k}(u_{i,k},v_{i,k})}{\partial u_{i,k}}\right|_{\boldsymbol{u}=\boldsymbol{v}} \\ &+\left(\boldsymbol{\nu}^\circ\cdot\left.\frac{\partial\boldsymbol{q}(\boldsymbol{z},\boldsymbol{v})}{\partial z_{j,l}}\right|_{\boldsymbol{z}=\boldsymbol{v}}\right)\times s_{i,k}(\boldsymbol{v})\times\left.\frac{\partial z_{j,l}(u_{j,l},v_{j,l})}{\partial u_{j,l}}\right|_{\boldsymbol{u}=\boldsymbol{v}} \end{aligned} \tag{A-30}$$

vanishes at an evolutionary singular trait $\boldsymbol{v}^*$ in the case of independent traits (i.e. when $s_{j,l}(\boldsymbol{v}^*)=0$ for any trait $(j,l)$).

##### A.3.2 Age-specific Hessian matrices

According to the second line of eq. (A-28), writing elements of Hessian matrices in terms of Hamiltonians requires expressions for $\partial^2 R_{mn}(\boldsymbol{z}_n,\boldsymbol{v})/\partial z_{i,k}\partial z_{j,l}$ in terms of Hamiltonians. Because these derivatives are non-zero only when both traits are expressed in same the parent class $n$ (i.e. $\partial^2 R_{mn}(\boldsymbol{z}_n,\boldsymbol{v})/\partial z_{i,k}\partial z_{j,l}\neq 0$ requires $i=j=n$), we can concentrate on finding expressions for $\partial^2 R_{mj}(\boldsymbol{z}_n,\boldsymbol{v})/\partial z_{j,k}\partial z_{j,l}$. To obtain these expressions, we start by differentiating eq. (A-15) with respect to traits $z_{j,k}$ and $z_{j,l}$. Recalling that mutant state values $\boldsymbol{x}_n(a)$ depend on trait expression form birth to age $a$, by virtue of the chain rule and the product rule we have

$$
\begin{aligned}
\left.\frac{\partial^2 R_{mj}(\boldsymbol{z}_j,\boldsymbol{v})}{\partial z_{j,k}\partial z_{j,l}}\right|_{\boldsymbol{z}=\boldsymbol{v}} = \int_0^\infty \Bigg[ & \left.\frac{\partial^2 H_{mj}(\boldsymbol{z}_j,\boldsymbol{x}_j^\circ(a),\boldsymbol{v})}{\partial z_{j,k}\partial z_{j,l}}\right|_{\boldsymbol{z}=\boldsymbol{v}} \\
& + \left.\frac{\partial^2 H_{mj}(\boldsymbol{z}_j,\boldsymbol{x}_j(a),\boldsymbol{v})}{\partial z_{j,k}\partial \boldsymbol{x}_j(a)}\right|_{\boldsymbol{z}=\boldsymbol{v}} \cdot \left.\frac{\partial \boldsymbol{x}_j(a)}{\partial z_{j,l}}\right|_{\boldsymbol{z}=\boldsymbol{v}} \\
& + \left.\frac{\partial^2 H_{mj}(\boldsymbol{z}_j,\boldsymbol{x}_j(a),\boldsymbol{v})}{\partial z_{j,l}\partial \boldsymbol{x}_j(a)}\right|_{\boldsymbol{z}=\boldsymbol{v}} \cdot \left.\frac{\partial \boldsymbol{x}_j(a)}{\partial z_{j,k}}\right|_{\boldsymbol{z}=\boldsymbol{v}} \\
& + \left( \left.\frac{\partial^2 H_{mj}(\boldsymbol{z}_j,\boldsymbol{x}_j(a),\boldsymbol{v})}{\partial \boldsymbol{x}_j(a)^2}\right|_{\boldsymbol{z}=\boldsymbol{v}} \left.\frac{\partial \boldsymbol{x}_j(a)}{\partial z_{j,k}}\right|_{\boldsymbol{z}=\boldsymbol{v}} \right) \cdot \left.\frac{\partial \boldsymbol{x}_j(a)}{\partial z_{j,l}}\right|_{\boldsymbol{z}=\boldsymbol{v}} \\
& + \left( \left.\frac{\partial H_{mj}(\boldsymbol{z}_j,\boldsymbol{x}_j(a),\boldsymbol{v})}{\partial \boldsymbol{x}_j(a)}\right|_{\boldsymbol{z}=\boldsymbol{v}} + \frac{\mathrm{d}\tilde{\boldsymbol{\lambda}}_{mj}(a)}{\mathrm{d}a} \right) \cdot \left.\frac{\partial^2 \boldsymbol{x}_j(a)}{\partial z_{j,k}\partial z_{j,l}}\right|_{\boldsymbol{z}=\boldsymbol{v}} \Bigg] \mathrm{d}a \\
& - \tilde{\boldsymbol{\lambda}}_{mj}(\infty) \cdot \left.\frac{\partial^2 \boldsymbol{x}_j(\infty)}{\partial z_{j,k}\partial z_{j,l}}\right|_{\boldsymbol{z}=\boldsymbol{v}} + \tilde{\boldsymbol{\lambda}}_{mj}(0) \cdot \left.\frac{\partial^2 \boldsymbol{x}_j(0)}{\partial z_{j,k}\partial z_{j,l}}\right|_{\boldsymbol{z}=\boldsymbol{v}},
\end{aligned}
\tag{A-31}
$$

where the second derivative of $H_{mj}$ with respect to a $\boldsymbol{x}_j(a)$ (fourth line) is a matrix with element $(k,l)$ given by $\partial^2 H_{mj}/\partial\tilde{x}_{j,k}(a)\partial\tilde{x}_{j,l}(a)$ (we use this notation thorough), that collects the second order effects of state values on all components of class-oriented individual fecundity. Note that similarly to eq. (A-16), we slightly abused notation in the first line of eq. (A-31), preemptively substituting state values $\boldsymbol{x}_j(a)$ with resident state values $\boldsymbol{x}_j^\circ(a)$, to make more explicit the fact that the derivative of the Hamiltonian with respect to mutant traits is a direct derivative. Derivatives of the Hamiltonian with respect to mutant traits in the second and third line of eq. (A-31) are also direct derivative, but there the same notational trick is impossible.

As initial state values $\boldsymbol{x}_j(0)$ are by assumption independent of trait values (i.e. their derivatives with respect to $z_{j,l}$ vanish), we can see from eq. (A-31) that if costates satisfy eq. (A-17), then

$$
\begin{aligned}
\left.\frac{\partial^2 R_{mj}(\boldsymbol{z}_j,\boldsymbol{v})}{\partial z_{j,k}\partial z_{j,l}}\right|_{\boldsymbol{z}=\boldsymbol{v}} = \int_0^\infty \Bigg[ & \left.\frac{\partial^2 H_{mj}(\boldsymbol{z}_j,\boldsymbol{x}_j^\circ(a),\boldsymbol{v})}{\partial z_{j,k}\partial z_{j,l}}\right|_{\boldsymbol{z}=\boldsymbol{v}} \\
& + \underbrace{\left.\frac{\partial^2 H_{mj}(\boldsymbol{z}_j,\boldsymbol{x}_j(a),\boldsymbol{v})}{\partial z_{j,k}\partial \boldsymbol{x}_j(a)}\right|_{\boldsymbol{z}=\boldsymbol{v}} \cdot \left.\frac{\partial \boldsymbol{x}_j(a)}{\partial z_{j,l}}\right|_{\boldsymbol{z}=\boldsymbol{v}}}_{\mathrm{A}} \\
& + \underbrace{\left.\frac{\partial^2 H_{mj}(\boldsymbol{z}_j,\boldsymbol{x}_j(a),\boldsymbol{v})}{\partial z_{j,l}\partial \boldsymbol{x}_j(a)}\right|_{\boldsymbol{z}=\boldsymbol{v}} \cdot \left.\frac{\partial \boldsymbol{x}_j(a)}{\partial z_{j,k}}\right|_{\boldsymbol{z}=\boldsymbol{v}}}_{\mathrm{B}} \\
& + \underbrace{\left( \left.\frac{\partial^2 H_{mj}(\boldsymbol{z}_j,\boldsymbol{x}_j(a),\boldsymbol{v})}{\partial \boldsymbol{x}_j(a)^2}\right|_{\boldsymbol{z}=\boldsymbol{v}} \left.\frac{\partial \boldsymbol{x}_j(a)}{\partial z_{j,k}}\right|_{\boldsymbol{z}=\boldsymbol{v}} \right) \cdot \left.\frac{\partial \boldsymbol{x}_j(a)}{\partial z_{j,l}}\right|_{\boldsymbol{z}=\boldsymbol{v}}}_{\mathrm{C}} \Bigg] \mathrm{d}a,
\end{aligned}
\tag{A-32}
$$

where labelled terms can be partitioned to distinguish selective effects acting through survivorship from those acting via internal states and written as

$$\begin{aligned}\mathrm{A} = & \left.\frac{\partial^2 H_{mj}(\boldsymbol{z}_j, l_j^{\circ}(a), \boldsymbol{x}_j(a), \boldsymbol{v})}{\partial z_{j,k}\partial \boldsymbol{x}_j(a)}\right|_{\boldsymbol{z}=\boldsymbol{v}} \cdot \left.\frac{\partial \boldsymbol{x}_j(a)}{\partial z_{j,l}}\right|_{\boldsymbol{z}=\boldsymbol{v}} \\ & + \left.\left(\frac{\partial^2 H_{mj}(\boldsymbol{z}_j, l_j(a), \boldsymbol{x}_j^{\circ}(a), \boldsymbol{v})}{\partial z_{j,k}\partial l_j(a)} \frac{\partial l_j(a)}{\partial z_{j,l}}\right)\right|_{\boldsymbol{z}=\boldsymbol{v}},\end{aligned} \tag{A-33}$$

$$\begin{aligned}\mathrm{B} = & \left.\frac{\partial^2 H_{mj}(\boldsymbol{z}_j, l_j^{\circ}(a), \boldsymbol{x}_j(a), \boldsymbol{v})}{\partial z_{j,l}\partial \boldsymbol{x}_j(a)}\right|_{\boldsymbol{z}=\boldsymbol{v}} \cdot \left.\frac{\partial \boldsymbol{x}_j(a)}{\partial z_{j,k}}\right|_{\boldsymbol{z}=\boldsymbol{v}} \\ & + \left.\left(\frac{\partial^2 H_{mj}(\boldsymbol{z}_j, l_j(a), \boldsymbol{x}_j^{\circ}(a), \boldsymbol{v})}{\partial z_{j,l}\partial l_j(a)} \frac{\partial l_j(a)}{\partial z_{j,k}}\right)\right|_{\boldsymbol{z}=\boldsymbol{v}},\end{aligned} \tag{A-34}$$

and

$$\begin{aligned}\mathrm{C} = & \left.\left(\frac{\partial^2 H_{mj}(\boldsymbol{z}_j, l_j^{\circ}(a), \boldsymbol{x}_j(a), \boldsymbol{v})}{\partial \boldsymbol{x}_j(a)^2} \frac{\partial \boldsymbol{x}_j(a)}{\partial z_{j,k}}\right)\right|_{\boldsymbol{z}=\boldsymbol{v}} \cdot \left.\frac{\partial \boldsymbol{x}_j(a)}{\partial z_{j,l}}\right|_{\boldsymbol{z}=\boldsymbol{v}} \\ & + \left.\left(\frac{\partial l_j(a)}{\partial z_{j,k}} \frac{\partial \boldsymbol{x}_j(a)}{\partial z_{j,l}} + \frac{\partial l_j(a)}{\partial z_{j,l}} \frac{\partial \boldsymbol{x}_j(a)}{\partial z_{j,k}}\right)\right|_{\boldsymbol{z}=\boldsymbol{v}} \cdot \left.\frac{\partial^2 H_{mj}(\boldsymbol{z}_j, l_j(a), \boldsymbol{x}_j(a), \boldsymbol{v})}{\partial l_j(a)\partial \boldsymbol{x}_j(a)}\right|_{\boldsymbol{z}=\boldsymbol{v}} \\ & + \left.\left(\underbrace{\frac{\partial^2 H_{mj}(\boldsymbol{z}_j, l_j(a), \boldsymbol{x}_j^{\circ}(a), \boldsymbol{v})}{\partial l_j(a)^2}}_{0} \frac{\partial l_j(a)}{\partial z_{j,k}} \frac{\partial l_j(a)}{\partial z_{j,l}}\right)\right|_{\boldsymbol{z}=\boldsymbol{v}}.\end{aligned} \tag{A-35}$$

where the last line is zero as the Hamiltonians depend only linearly on survivorship.

We can now use each of the terms that constitute $\partial^2 R_{mj}(\boldsymbol{z}_j, \boldsymbol{v})/\partial z_{j,k}\partial z_{j,l} \mid_{\boldsymbol{z}=\boldsymbol{v}}$, listed in eqs. (A-32)–(A-35), to obtain eqs. (16a)–(16e) of the main text. Substituting the term from the first line of eq. (A-32) into the first line of eq. (A-28) for the case where $i = j$ (recall that $\partial^2 R_{mn}(\boldsymbol{z}_n, \boldsymbol{v})/\partial z_{i,k}\partial z_{j,l} \neq 0$ requires $i = j = n$) gives

$$\begin{aligned}h_{jj,kl}^{\mathrm{zz}}(\boldsymbol{v}) = & \int_0^{\infty} \left[\sum_{m=1}^{n_{\mathrm{c}}} \nu_m^{\circ} \gamma_{mj} \left.\frac{\partial^2 H_{mj}(\boldsymbol{z}_j, l_j^{\circ}(a), \boldsymbol{x}_j^{\circ}(a), \boldsymbol{v})}{\partial z_{j,k}\partial z_{j,l}}\right|_{\boldsymbol{z}=\boldsymbol{v}} q_j^{\circ}\right] \mathrm{d}a \\ & \times \left.\frac{\partial z_{i,k}(u_{i,k}, v_{i,k})}{\partial u_{i,k}}\right|_{\boldsymbol{u}=\boldsymbol{v}} \times \left.\frac{\partial z_{j,l}(u_{j,l}, v_{j,l})}{\partial u_{j,l}}\right|_{\boldsymbol{u}=\boldsymbol{v}},\end{aligned} \tag{A-36}$$

where the integrand corresponds to eq. (16a) of the main text. Similarly, plugging the term of eq. (A-32) corresponding to the first line of eq. (A-35) into the first line of eq. (A-28) gives

$$\begin{aligned}h_{jj,kl}^{\mathrm{xx}}(\boldsymbol{v}) = & \int_0^{\infty} \left[\sum_{m=1}^{n_{\mathrm{c}}} \nu_m^{\circ} \gamma_{mj} \left(\left.\left(\frac{\partial^2 H_{mj}(\boldsymbol{z}_j, l_j^{\circ}(a), \boldsymbol{x}_j(a), \boldsymbol{v})}{\partial \boldsymbol{x}_j(a)^2} \frac{\partial \boldsymbol{x}_j(a)}{\partial z_{j,k}}\right)\right|_{\boldsymbol{z}=\boldsymbol{v}} \cdot \left.\frac{\partial \boldsymbol{x}_j(a)}{\partial z_{j,l}}\right|_{\boldsymbol{z}=\boldsymbol{v}}\right) q_j^{\circ}\right] \mathrm{d}a \\ & \times \left.\frac{\partial z_{i,k}(u_{i,k}, v_{i,k})}{\partial u_{i,k}}\right|_{\boldsymbol{u}=\boldsymbol{v}} \times \left.\frac{\partial z_{j,l}(u_{j,l}, v_{j,l})}{\partial u_{j,l}}\right|_{\boldsymbol{u}=\boldsymbol{v}},\end{aligned} \tag{A-37}$$

where the integrand corresponds to eq. (16b) of the main text in matrix form. Likewise, using the terms from the first line of eq. (A-33) and the first line of eq. (A-34) in the first line of eq. (A-28) gives

$$\begin{aligned} h^{\mathrm{zx}}_{jj,kl}(\boldsymbol{v}) = \int_0^\infty \Bigg[ \sum_{m=1}^{n_\mathrm{c}} \nu^\circ_m \gamma_{mj} \Bigg( & \left.\frac{\partial^2 H_{mj}(\boldsymbol{z}_j, l^\circ_j(a), \boldsymbol{x}_j(a), \boldsymbol{v})}{\partial z_{j,k} \partial \boldsymbol{x}_j(a)}\right|_{\boldsymbol{z}=\boldsymbol{v}} \cdot \left.\frac{\partial \boldsymbol{x}_j(a)}{\partial z_{j,l}}\right|_{\boldsymbol{z}=\boldsymbol{v}} \\ & + \left.\frac{\partial^2 H_{mj}(\boldsymbol{z}_j, l^\circ_j(a), \boldsymbol{x}_j(a), \boldsymbol{v})}{\partial z_{j,l} \partial \boldsymbol{x}_j(a)}\right|_{\boldsymbol{z}=\boldsymbol{v}} \cdot \left.\frac{\partial \boldsymbol{x}_j(a)}{\partial z_{j,k}}\right|_{\boldsymbol{z}=\boldsymbol{v}} \Bigg) q^\circ_j \Bigg] \mathrm{d}a \\ & \times \left.\frac{\partial z_{i,k}(u_{i,k}, v_{i,k})}{\partial u_{i,k}}\right|_{\boldsymbol{u}=\boldsymbol{v}} \times \left.\frac{\partial z_{j,l}(u_{j,l}, v_{j,l})}{\partial u_{j,l}}\right|_{\boldsymbol{u}=\boldsymbol{v}}, \end{aligned} \tag{A-38}$$

where the integrand corresponds to eq. (16c) of the main text in vector form. Now using, the terms from the second line of eq. (A-33) and the second line of eq. (A-34) in the first line of eq. (A-28) gives

$$\begin{aligned} h^{\mathrm{zl}}_{jj,kl}(\boldsymbol{v}) = \int_0^\infty \Bigg[ \sum_{m=1}^{n_\mathrm{c}} \nu^\circ_m \gamma_{mj} \Bigg( & \left.\left( \frac{\partial^2 H_{mj}(\boldsymbol{z}_j, l_j(a), \boldsymbol{x}^\circ_j(a), \boldsymbol{v})}{\partial z_{j,k} \partial l_j(a)} \frac{\partial l_j(a)}{\partial z_{j,l}} \right)\right|_{\boldsymbol{z}=\boldsymbol{v}} \\ & + \left.\left( \frac{\partial^2 H_{mj}(\boldsymbol{z}_j, l_j(a), \boldsymbol{x}^\circ_j(a), \boldsymbol{v})}{\partial z_{j,l} \partial l_j(a)} \frac{\partial l_j(a)}{\partial z_{j,k}} \right)\right|_{\boldsymbol{z}=\boldsymbol{v}} \Bigg) q^\circ_j \Bigg] \mathrm{d}a \\ & \times \left.\frac{\partial z_{i,k}(u_{i,k}, v_{i,k})}{\partial u_{i,k}}\right|_{\boldsymbol{u}=\boldsymbol{v}} \times \left.\frac{\partial z_{j,l}(u_{j,l}, v_{j,l})}{\partial u_{j,l}}\right|_{\boldsymbol{u}=\boldsymbol{v}}, \end{aligned} \tag{A-39}$$

where the integrand corresponds to eq. (16d) of the main text. Finally, using the term from the second line of eq. (A-35) in the first line of eq. (A-28) gives

$$\begin{aligned} h^{\mathrm{lx}}_{jj,kl}(\boldsymbol{v}) = \int_0^\infty \Bigg[ \sum_{m=1}^{n_\mathrm{c}} \nu^\circ_m \gamma_{mj} \Bigg( & \left.\left( \frac{\partial l_j(a)}{\partial z_{j,k}} \frac{\partial \boldsymbol{x}_j(a)}{\partial z_{j,l}} + \frac{\partial l_j(a)}{\partial z_{j,l}} \frac{\partial \boldsymbol{x}_j(a)}{\partial z_{j,k}} \right)\right|_{\boldsymbol{z}=\boldsymbol{v}} \\ & \cdot \left.\frac{\partial^2 H_{mj}(\boldsymbol{z}_j, l_j(a), \boldsymbol{x}_j(a), \boldsymbol{v})}{\partial l_j(a) \partial \boldsymbol{x}_j(a)}\right|_{\boldsymbol{z}=\boldsymbol{v}} \Bigg) q^\circ_j \Bigg] \mathrm{d}a \\ & \times \left.\frac{\partial z_{i,k}(u_{i,k}, v_{i,k})}{\partial u_{i,k}}\right|_{\boldsymbol{u}=\boldsymbol{v}} \times \left.\frac{\partial z_{j,l}(u_{j,l}, v_{j,l})}{\partial u_{j,l}}\right|_{\boldsymbol{u}=\boldsymbol{v}}, \end{aligned} \tag{A-40}$$

where the integrand corresponds to eq. (16e) of the main text in vector form.

Next, we obtain integral representations for eqs. (A-29) and (A-30) (the last two terms of eq. (A-28)). Plugging eq. (A-18) into eq. (A-29) gives

$$\begin{aligned} h^{\mathrm{zq}}_{ij,kl}(\boldsymbol{v}) = & \left[ \int_0^\infty \sum_{m=1}^{n_\mathrm{c}} \nu^\circ_m \left.\left( \gamma_{mi} \frac{\partial H_{mi}(\boldsymbol{z}_i, \boldsymbol{v})}{\partial z_{i,k}} \frac{\partial q_i(\boldsymbol{z}, \boldsymbol{v})}{\partial z_{j,l}} + \gamma_{mj} \frac{\partial H_{mj}(\boldsymbol{z}_j, \boldsymbol{v})}{\partial z_{j,l}} \frac{\partial q_j(\boldsymbol{z}, \boldsymbol{v})}{\partial z_{i,k}} \right)\right|_{\boldsymbol{z}=\boldsymbol{v}} \mathrm{d}a \right] \\ & \times \left.\frac{\partial z_{i,k}(u_{i,k}, v_{i,k})}{\partial u_{i,k}}\right|_{\boldsymbol{u}=\boldsymbol{v}} \times \left.\frac{\partial z_{j,l}(u_{j,l}, v_{j,l})}{\partial u_{j,l}}\right|_{\boldsymbol{u}=\boldsymbol{v}}, \end{aligned} \tag{A-41}$$

where the integrand corresponds to eq. (16f) of the main text in vector form. In turn, plugging eq. (8)

into eq. (A-30) gives

$$h^{\mathrm{q}}_{ij,kl}(\boldsymbol{v}) = \left[\int_0^\infty \left(\boldsymbol{\nu}^\circ \cdot \left.\frac{\partial \boldsymbol{q}(\boldsymbol{z},\boldsymbol{v})}{\partial z_{i,k}}\right|_{\boldsymbol{z}=\boldsymbol{v}}\right) \times \hat{s}_{j,l}(a,\boldsymbol{v}) + \left(\boldsymbol{\nu}^\circ \cdot \left.\frac{\partial \boldsymbol{q}(\boldsymbol{z},\boldsymbol{v})}{\partial z_{j,l}}\right|_{\boldsymbol{z}=\boldsymbol{v}}\right) \times \hat{s}_{i,k}(a,\boldsymbol{v})\,\mathrm{d}a\right] \\ \times \left.\frac{\partial z_{i,k}(u_{i,k},v_{i,k})}{\partial u_{i,k}}\right|_{\boldsymbol{u}=\boldsymbol{v}} \times \left.\frac{\partial z_{j,l}(u_{j,l},v_{j,l})}{\partial u_{j,l}}\right|_{\boldsymbol{u}=\boldsymbol{v}}. \tag{A-42}$$

Besides derivatives of the Hamiltonians, eqs. (A-41) and (A-42) show that computing Hessian matrices requires computing state perturbations (i.e. $\partial l_n(a)/\partial z_{j,l}$ and $\partial \boldsymbol{x}_n(a)/\partial z_{j,l}$) as well as eigenvector perturbations (i.e. $\partial \boldsymbol{q}(\boldsymbol{z},\boldsymbol{v})/\partial z_{j,l}$) with respect to any trait $z_{j,l}$. We give methods to compute these quantities in the next section.

#### A.4 State and eigenvector perturbations

##### A.4.1 State perturbation

Denoting $\boldsymbol{x}_{j,0} = \boldsymbol{x}_j(0) = (1, \boldsymbol{x}_{j,0})$ the concatenation of initial values for survivorship and internal states, we start by recognizing that by the fundamental theorem of calculus, mutant states at age $a$ in class $j$ can be written as

$$\boldsymbol{x}_j(a) = \boldsymbol{x}_{j,0} + \int_0^a \tilde{\boldsymbol{g}}_j(\boldsymbol{z}_j, \boldsymbol{x}_j(\tau), \boldsymbol{v})\mathrm{d}\tau. \tag{A-43}$$

Under our assumption that initial state values $\boldsymbol{x}_{n,0}$ are independent of traits, using the chain rule and remembering that state values depend on trait values (except the initial values), eq. (A-43) implies that

$$\left.\frac{\partial \boldsymbol{x}_j(a)}{\partial z_{j,l}}\right|_{\boldsymbol{z}=\boldsymbol{v}} = \int_0^a \left[\left.\frac{\partial \tilde{\boldsymbol{g}}_j(\boldsymbol{z}_j, \boldsymbol{x}_j(\tau), \boldsymbol{v})}{\partial \boldsymbol{x}_j(\tau)}\right|_{\boldsymbol{z}=\boldsymbol{v}} \left.\frac{\partial \boldsymbol{x}_j(\tau)}{\partial z_{j,l}}\right|_{\boldsymbol{z}=\boldsymbol{v}} + \left.\frac{\partial \tilde{\boldsymbol{g}}_j(\boldsymbol{z}_j, \boldsymbol{x}^\circ_j(\tau), \boldsymbol{v})}{\partial z_{j,l}}\right|_{\boldsymbol{z}=\boldsymbol{v}}\right] \mathrm{d}\tau. \tag{A-44}$$

Recalling that $\boldsymbol{x}_j(a) = (l_j(a), \boldsymbol{x}_j(a))$ and $\tilde{\boldsymbol{g}}_j(\boldsymbol{z}_j, \boldsymbol{x}_j(\tau), \boldsymbol{v}) = \big(-\mu_j(\boldsymbol{z}_j, \boldsymbol{x}_j(a), \boldsymbol{v})l_j(a), \boldsymbol{g}_j(\boldsymbol{z}_j, \boldsymbol{x}_j(\tau), \boldsymbol{v})\big)$, we can see that

$$\begin{aligned} \left.\frac{\partial \tilde{\boldsymbol{g}}_j(\boldsymbol{z}_j, \boldsymbol{x}_j(\tau), \boldsymbol{v})}{\partial \boldsymbol{x}_j(\tau)}\right|_{\boldsymbol{z}=\boldsymbol{v}} &= \begin{bmatrix} -\left.\frac{\partial \mu_j(\boldsymbol{z}_j, \boldsymbol{x}^\circ_j(\tau), \boldsymbol{v})l_j(\tau)}{\partial l_j(\tau)}\right|_{\boldsymbol{z}=\boldsymbol{v}} & -\left.\frac{\partial \mu_j(\boldsymbol{z}_j, \boldsymbol{x}_j(\tau), \boldsymbol{v})l^\circ_j(\tau)}{\partial \boldsymbol{x}_j(\tau)}\right|_{\boldsymbol{z}=\boldsymbol{v}} \\ \left.\frac{\partial \boldsymbol{g}_j(\boldsymbol{z}_j, \boldsymbol{x}^\circ_j(\tau), \boldsymbol{v})}{\partial l_j(\tau)}\right|_{\boldsymbol{z}=\boldsymbol{v}} & \left.\frac{\partial \boldsymbol{g}_j(\boldsymbol{z}_j, \boldsymbol{x}_j(\tau), \boldsymbol{v})}{\partial \boldsymbol{x}_j(\tau)}\right|_{\boldsymbol{z}=\boldsymbol{v}} \end{bmatrix} \\ &= \begin{bmatrix} -\mu_j(\boldsymbol{v}_j, \boldsymbol{x}^\circ_j(\tau), \boldsymbol{v}) & -l^\circ_j(\tau) \left.\frac{\partial \mu_j(\boldsymbol{z}_j, \boldsymbol{x}_j(\tau), \boldsymbol{v})}{\partial \boldsymbol{x}_j(\tau)}\right|_{\boldsymbol{z}=\boldsymbol{v}} \\ 0 & \left.\frac{\partial \boldsymbol{g}_j(\boldsymbol{z}_j, \boldsymbol{x}_j(\tau), \boldsymbol{v})}{\partial \boldsymbol{x}_j(\tau)}\right|_{\boldsymbol{z}=\boldsymbol{v}} \end{bmatrix} \end{aligned} \tag{A-45}$$

is a matrix collecting the sensitivity of the rates of state change in class $n$ to a change in state values. With this, differentiating both sides of eq. (A-44) with respect to age $a$ yields the system of ODE's

$$\begin{aligned}\frac{\mathrm{d}}{\mathrm{d}a}\left[\left.\frac{\partial l_j(a)}{\partial z_{j,l}}\right|_{\boldsymbol{z}=\boldsymbol{v}}\right] &= -l_j^{\circ}(a)\left.\frac{\partial \mu_j(\boldsymbol{z}_j,\boldsymbol{x}_j(a),\boldsymbol{v})}{\partial \boldsymbol{x}_j(a)}\right|_{\boldsymbol{z}=\boldsymbol{v}}\cdot\frac{\partial \boldsymbol{x}_j(a)}{\partial z_{j,l}} - l_j^{\circ}(a)\left.\frac{\partial \mu_j(\boldsymbol{z}_j,\boldsymbol{x}_j^{\circ}(a),\boldsymbol{v})}{\partial z_{j,l}}\right|_{\boldsymbol{z}=\boldsymbol{v}} \\ &\quad - \mu_j(\boldsymbol{v}_j,\boldsymbol{x}_j^{\circ}(a),\boldsymbol{v})\left.\frac{\partial l_j(a)}{\partial z_{j,l}}\right|_{\boldsymbol{z}=\boldsymbol{v}} \quad \text{with i.c.} \quad \left.\frac{\partial l_j(0)}{\partial z_{j,l}}\right|_{\boldsymbol{z}=\boldsymbol{v}} = 0, \\ \frac{\mathrm{d}}{\mathrm{d}a}\left[\left.\frac{\partial \boldsymbol{x}_j(a)}{\partial z_{j,l}}\right|_{\boldsymbol{z}=\boldsymbol{v}}\right] &= \left.\frac{\partial \boldsymbol{g}_j(\boldsymbol{z}_j,\boldsymbol{x}_j(a),\boldsymbol{v})}{\partial \boldsymbol{x}_j(a)}\right|_{\boldsymbol{z}=\boldsymbol{v}}\left.\frac{\partial \boldsymbol{x}_j(a)}{\partial z_{j,l}}\right|_{\boldsymbol{z}=\boldsymbol{v}} \\ &\quad + \left.\frac{\partial \boldsymbol{g}_j(\boldsymbol{z}_j,\boldsymbol{x}_j^{\circ}(a),\boldsymbol{v})}{\partial z_{j,l}}\right|_{\boldsymbol{z}=\boldsymbol{v}} \quad \text{with i.c.} \quad \left.\frac{\partial \boldsymbol{x}_j(0)}{\partial z_{j,l}}\right|_{\boldsymbol{z}=\boldsymbol{v}} = \mathbf{0}.\end{aligned} \tag{A-46}$$

We can now study the solution to eq. (A-46). According to the generalized Cauchy formula (Weber 2011, p.72), an inhomogeneous linear system of the form $\mathrm{d}\mathbf{y}(a)/\mathrm{d}a = \mathbf{b}(a) + \mathbf{A}(a)\mathbf{y}(a)$, where $\mathbf{b}(a)$ and $\mathbf{A}(a)$ are an arbitrary vector and matrix, admits a solution of the form

$$\mathbf{y}(a) = \boldsymbol{\Psi}(a,\zeta)\mathbf{y}(\zeta) + \int_{\zeta}^{a}\boldsymbol{\Psi}(a,\tau)\mathbf{b}(\tau)\mathrm{d}\tau, \tag{A-47}$$

where $\boldsymbol{\Psi}(a,\tau)$ is the so-called fundamental matrix, which satisfies

$$\frac{\mathrm{d}\boldsymbol{\Psi}(a,\tau)}{\mathrm{d}a} = \mathbf{A}(a)\boldsymbol{\Psi}(a,\tau) \text{ with i.c. } \boldsymbol{\Psi}(\tau,\tau) = \mathbf{I} \tag{A-48}$$

with $\mathbf{I}$ denoting the $n_{\mathrm{x}} \times n_{\mathrm{x}}$ identity matrix. Then, using eq. (A-47) with $\zeta = 0$, holding internal state perturbations as a known quantity and assuming that traits have no effect on survivorship at birth (i.e. $\partial l_j(0)/\partial z_{j,l} = 0$), survival perturbations admit a solution of the form

$$\begin{aligned}\left.\frac{\partial l_j(a)}{\partial z_{j,l}}\right|_{\boldsymbol{z}=\boldsymbol{v}} &= \int_0^a \frac{l_j^{\circ}(a)}{l_j^{\circ}(\tau)} \\ &\quad \times\left(-l_j^{\circ}(\tau)\left.\frac{\partial \mu_j(\boldsymbol{z}_j,\boldsymbol{x}_j(\tau),\boldsymbol{v})}{\partial \boldsymbol{x}_j(\tau)}\right|_{\boldsymbol{z}=\boldsymbol{v}}\cdot\left.\frac{\partial \boldsymbol{x}_j(\tau)}{\partial z_{j,l}}\right|_{\boldsymbol{z}=\boldsymbol{v}} - l_j^{\circ}(\tau)\left.\frac{\partial \mu_j(\boldsymbol{z}_j,\boldsymbol{x}_j^{\circ}(\tau),\boldsymbol{v})}{\partial z_{j,l}}\right|_{\boldsymbol{z}=\boldsymbol{v}}\right)\mathrm{d}\tau \\ &= -l_j^{\circ}(a)\int_0^a\left(\left.\frac{\partial \mu_j(\boldsymbol{z}_j,\boldsymbol{x}_j(\tau),\boldsymbol{v})}{\partial \boldsymbol{x}_j(\tau)}\right|_{\boldsymbol{z}=\boldsymbol{v}}\cdot\left.\frac{\partial \boldsymbol{x}_j(\tau)}{\partial z_{j,l}}\right|_{\boldsymbol{z}=\boldsymbol{v}} + \left.\frac{\partial \mu_j(\boldsymbol{z}_j,\boldsymbol{x}_j^{\circ}(\tau),\boldsymbol{v})}{\partial z_{j,l}}\right|_{\boldsymbol{z}=\boldsymbol{v}}\right)\mathrm{d}\tau,\end{aligned} \tag{A-49}$$

where the ratio $l_j^{\circ}(a)/l_j^{\circ}(\tau) = \exp\left(-\int_{\tau}^{a}\mu_j(\boldsymbol{v}_j,\boldsymbol{x}_j^{\circ}(s),\boldsymbol{v})\mathrm{d}s\right)$ here is the one-dimensional equivalent of the fundamental matrix, obtained as the solution $\Psi_j(a,\tau)$ to

$$\frac{\mathrm{d}\Psi_j(a,\tau)}{\mathrm{d}a} = -\mu_j(\boldsymbol{v}_j,\boldsymbol{x}_j^{\circ}(a),\boldsymbol{v})\Psi_j(a,\tau) \text{ with i.c. } \Psi_j(\tau,\tau) = 1, \tag{A-50}$$

obtained by applying eq. (A-48). In addition, state perturbations admit a solution of the form

$$\left.\frac{\partial \boldsymbol{x}_j(a)}{\partial z_{j,l}}\right|_{\boldsymbol{z}=\boldsymbol{v}} = \int_0^a \boldsymbol{\Psi}_j(a,\tau)\left.\frac{\partial \boldsymbol{g}_j(\boldsymbol{z}_j,\boldsymbol{x}_j^{\circ}(\tau),\boldsymbol{v})}{\partial z_{j,l}}\right|_{\boldsymbol{z}=\boldsymbol{v}}\mathrm{d}\tau, \tag{A-51}$$

with $\boldsymbol{\Psi}_j$ the $(n_\mathrm{x} \times n_\mathrm{x})$ fundamental matrix associated to internal states perturbations satisfying

$$\frac{\mathrm{d}\boldsymbol{\Psi}_j(a,\tau)}{\mathrm{d}a} = \left.\frac{\partial \boldsymbol{g}_j(\boldsymbol{z}_j, \boldsymbol{x}_j(a), \boldsymbol{v})}{\partial \boldsymbol{x}_j(a)}\right|_{\boldsymbol{z}=\boldsymbol{v}} \boldsymbol{\Psi}_j(a,\tau) \text{ with i.c. } \boldsymbol{\Psi}_j(\tau,\tau) = \mathbf{I}. \tag{A-52}$$

Because the coefficient matrix $\partial \boldsymbol{g}_j(\boldsymbol{z}_j, \boldsymbol{x}_j(a), \boldsymbol{v})/\partial \boldsymbol{x}_j(a)$ is not constant in general, there is no general explicit expression for $\boldsymbol{\Psi}_j$, and thus none for the state perturbations either. In the case where the coefficient matrix is constant, the fundamental matrix can be expressed as

$$\boldsymbol{\Psi}_j(a,\tau) = \exp\left( (a-\tau) \left.\frac{\partial \boldsymbol{g}_j(\boldsymbol{z}_j, \boldsymbol{x}_j(a), \boldsymbol{v})}{\partial \boldsymbol{x}_j(a)}\right|_{\boldsymbol{z}=\boldsymbol{v}} \right), \tag{A-53}$$

and thus in this case

$$\left.\frac{\partial \boldsymbol{x}_j(a)}{\partial z_{j,l}}\right|_{\boldsymbol{z}=\boldsymbol{v}} = \int_0^a \exp\left( (a-\tau) \left.\frac{\partial \boldsymbol{g}_j(\boldsymbol{z}_j, \boldsymbol{x}_j(a), \boldsymbol{v})}{\partial \boldsymbol{x}_j(a)}\right|_{\boldsymbol{z}=\boldsymbol{v}} \right) \left.\frac{\partial \boldsymbol{g}_j(\boldsymbol{z}_j, \boldsymbol{x}_j^\circ(\tau), \boldsymbol{v})}{\partial z_{j,l}}\right|_{\boldsymbol{z}=\boldsymbol{v}} \mathrm{d}\tau. \tag{A-54}$$

For a non-constant coefficient matrix numerical integration is usually required, but we note that arbitrarily close estimates of the fundamental matrix can be obtained by truncation of the Peano-Baker series (Weber 2011). Whether or not internal state perturbations have an explicit formula, once they are obtained they can be plugged into eq. (A-49) to compute survivorship perturbations.

##### A.4.2 Right-eigenvector perturbation

To obtain an explicit expression for derivatives of the leading right-eigenvector of the next-generation matrix $\mathbf{R}(\boldsymbol{z}, \boldsymbol{v})$ (given by eq. A-1), we start by differentiating eq. (A-2) with respect to $z_{j,l}$. Using the product rule gives

$$\left.\frac{\partial R_0(\boldsymbol{z}, \boldsymbol{v})}{\partial z_{j,l}}\right|_{\boldsymbol{z}=\boldsymbol{v}} \boldsymbol{q}^\circ + \left.\frac{\partial \boldsymbol{q}(\boldsymbol{z}, \boldsymbol{v})}{\partial z_{j,l}}\right|_{\boldsymbol{z}=\boldsymbol{v}} = \left.\frac{\partial \mathbf{R}(\boldsymbol{z}, \boldsymbol{v})}{\partial z_{j,l}}\right|_{\boldsymbol{z}=\boldsymbol{v}} \boldsymbol{q}^\circ + \mathbf{R}^\circ \left.\frac{\partial \boldsymbol{q}(\boldsymbol{z}, \boldsymbol{v})}{\partial z_{j,l}}\right|_{\boldsymbol{z}=\boldsymbol{v}}, \tag{A-55}$$

where we used the fact that $R_0(\boldsymbol{v}, \boldsymbol{v}) = 1$, and where $\partial \mathbf{R}(\boldsymbol{z}, \boldsymbol{v})/\partial z_{j,l}|_{\boldsymbol{u}=\boldsymbol{v}}$ is a matrix with element $(m, n)$ given by

$$\gamma_{mn} \left.\frac{\partial R_{mn}(\boldsymbol{z}, \boldsymbol{v})}{\partial z_{j,l}}\right|_{\boldsymbol{z}=\boldsymbol{v}} = \gamma_{mn} \int_0^\infty \left.\frac{\partial H_{mn}(\boldsymbol{z}_n, \boldsymbol{x}_n(a), \boldsymbol{v})}{\partial z_{j,l}}\right|_{\boldsymbol{z}=\boldsymbol{v}} \mathrm{d}a, \tag{A-56}$$

according to eq. (A-18).

After rearrangement, this leads to

$$(\mathbf{R}^\circ - \mathbf{I}) \left.\frac{\partial \boldsymbol{q}(\boldsymbol{z}, \boldsymbol{v})}{\partial z_{j,l}}\right|_{\boldsymbol{z}=\boldsymbol{v}} = \left( \left.\frac{\partial R_0(\boldsymbol{z}, \boldsymbol{v})}{\partial z_{j,l}}\right|_{\boldsymbol{z}=\boldsymbol{v}} \mathbf{I} - \left.\frac{\partial \mathbf{R}(\boldsymbol{z}, \boldsymbol{v})}{\partial z_{j,l}}\right|_{\boldsymbol{z}=\boldsymbol{v}} \right) \boldsymbol{q}^\circ. \tag{A-57}$$

The matrix $(\mathbf{R}^\circ - \mathbf{I})$ cannot be inverted, as $\det(\mathbf{R}^\circ - \mathbf{I}) = 0$ by the fact that the leading eigenvalue of $\mathbf{R}^\circ$ is $R_0(\boldsymbol{v}, \boldsymbol{v}) = 1$. To solve for the required derivatives, we associate one more equation to the linear system (A-57) that will make it invertible. To that end, let us introduce the $(n_c+1)\times n_c$ matrix

$$\check{\mathbf{R}}^{\circ} = \begin{bmatrix} (\mathbf{R}^{\circ}-\mathbf{I}) \\ \mathbf{1}^{(1\times n_c)} \end{bmatrix}, \tag{A-58}$$

where $\mathbf{1}^{(1\times n_c)}$ denotes a row vector of ones of size $n_c$, as well as the column vector of size $(n_c+1)$

$$\check{\boldsymbol{p}}_0 = \left( \left( \left.\frac{\partial R_0(\boldsymbol{z},\boldsymbol{v})}{\partial z_{j,l}}\right|_{\boldsymbol{z}=\boldsymbol{v}} \mathbf{I} - \left.\frac{\partial \mathbf{R}(\boldsymbol{z},\boldsymbol{v})}{\partial z_{j,l}}\right|_{\boldsymbol{z}=\boldsymbol{v}} \right) \boldsymbol{q}^{\circ}, 0 \right). \tag{A-59}$$

Using these expressions along eq. (A-57) and the constraint

$$\mathbf{1}^{(1\times n_c)} \cdot \left.\frac{\partial \boldsymbol{q}(\boldsymbol{z},\boldsymbol{v})}{\partial z_{j,l}}\right|_{\boldsymbol{z}=\boldsymbol{v}} = 0, \tag{A-60}$$

which follows from the conservation of total frequency in a lineage, we obtain that

$$\check{\mathbf{R}}^{\circ} \left.\frac{\partial \boldsymbol{q}(\boldsymbol{z},\boldsymbol{v})}{\partial z_{j,l}}\right|_{\boldsymbol{z}=\boldsymbol{v}} = \check{\boldsymbol{p}}_0. \tag{A-61}$$

Because the matrix $\check{\mathbf{R}}^{\circ}$ has rank $n_c$ (Fox & Kapoor 1968; Nelson 1976), it admits a left-inverse $\check{\mathbf{R}}^{\circ}_{\mathrm{L}} = (\check{\mathbf{R}}^{\circ\mathsf{T}}\check{\mathbf{R}}^{\circ})^{-1}\check{\mathbf{R}}^{\circ\mathsf{T}}$ such that $\check{\mathbf{R}}^{\circ}_{\mathrm{L}}\check{\mathbf{R}}^{\circ} = \mathbf{I}$ (where the minus one superscript refers to the standard inverse; Rao & Mitra (1972), p.19). Thus,

$$\left.\frac{\partial \boldsymbol{q}(\boldsymbol{z},\boldsymbol{v})}{\partial z_{j,l}}\right|_{\boldsymbol{z}=\boldsymbol{v}} = \left(\check{\mathbf{R}}^{\circ\mathsf{T}}\check{\mathbf{R}}^{\circ}\right)^{-1} \check{\mathbf{R}}^{\circ\mathsf{T}}\check{\boldsymbol{p}}_0. \tag{A-62}$$

Plugging eqs. (A-58) and (A-59) into eq. (A-62) produces

$$\begin{aligned}
\left.\frac{\partial \boldsymbol{q}(\boldsymbol{z},\boldsymbol{v})}{\partial z_{j,l}}\right|_{\boldsymbol{z}=\boldsymbol{v}} &= \left( \begin{bmatrix} (\mathbf{R}^{\circ}-\mathbf{I}) \\ \mathbf{1}^{(1\times n_c)} \end{bmatrix}^{\mathsf{T}} \begin{bmatrix} (\mathbf{R}^{\circ}-\mathbf{I}) \\ \mathbf{1}^{(1\times n_c)} \end{bmatrix} \right)^{-1} \\
&\quad \times \begin{bmatrix} (\mathbf{R}^{\circ}-\mathbf{I}) \\ \mathbf{1}^{(1\times n_c)} \end{bmatrix}^{\mathsf{T}} \left( \left( \left.\frac{\partial R_0(\boldsymbol{z},\boldsymbol{v})}{\partial z_{j,l}}\right|_{\boldsymbol{z}=\boldsymbol{v}} \mathbf{I} - \left.\frac{\partial \mathbf{R}(\boldsymbol{z},\boldsymbol{v})}{\partial z_{j,l}}\right|_{\boldsymbol{z}=\boldsymbol{v}} \right) \boldsymbol{q}^{\circ}, 0 \right) \\
&= \left( (\mathbf{R}^{\circ}-\mathbf{I})^{\mathsf{T}}(\mathbf{R}^{\circ}-\mathbf{I}) + \mathbf{1}^{(n_c\times 1)} \times \mathbf{1}^{(1\times n_c)} \right)^{-1} \\
&\quad \times \left( (\mathbf{R}^{\circ}-\mathbf{I})^{\mathsf{T}} \left( \left.\frac{\partial R_0(\boldsymbol{z},\boldsymbol{v})}{\partial z_{j,l}}\right|_{\boldsymbol{z}=\boldsymbol{v}} \mathbf{I} - \left.\frac{\partial \mathbf{R}(\boldsymbol{z},\boldsymbol{v})}{\partial z_{j,l}}\right|_{\boldsymbol{z}=\boldsymbol{v}} \right) \boldsymbol{q}^{\circ} + \mathbf{1}^{(n_c\times 1)} \times 0 \right),
\end{aligned} \tag{A-63}$$

where $\mathbf{1}^{(n_c\times n_c)}$ is a matrix of ones of dimensions $(n_c \times n_c)$. Using the fact that $\left.\frac{\partial R_0(\boldsymbol{z},\boldsymbol{v})}{\partial z_{j,l}}\right|_{\boldsymbol{z}=\boldsymbol{v}} = \left( \left.\frac{\partial z_{j,l}(u_{j,l},v_{j,l})}{\partial u_{j,l}}\right|_{\boldsymbol{u}=\boldsymbol{v}} \right)^{-1} s_{j,l}(\boldsymbol{v})$ (according to eq. 3b), we finally have

$$\begin{aligned}
\left.\frac{\partial \boldsymbol{q}(\boldsymbol{z},\boldsymbol{v})}{\partial z_{j,l}}\right|_{\boldsymbol{u}=\boldsymbol{v}} &= \left( (\mathbf{R}^{\circ\mathsf{T}}-\mathbf{I})(\mathbf{R}^{\circ}-\mathbf{I}) + \mathbf{1}^{(n_c\times n_c)} \right)^{-1} \\
&\quad \times (\mathbf{R}^{\circ\mathsf{T}}-\mathbf{I}) \left( \left( \left.\frac{\partial z_{j,l}(u_{j,l},v_{j,l})}{\partial u_{j,l}}\right|_{\boldsymbol{u}=\boldsymbol{v}} \right)^{-1} s_{j,l}(\boldsymbol{v})\mathbf{I} - \left.\frac{\partial \mathbf{R}(\boldsymbol{z},\boldsymbol{v})}{\partial z_{j,l}}\right|_{\boldsymbol{z}=\boldsymbol{v}} \right) \boldsymbol{q}^{\circ},
\end{aligned} \tag{A-64}$$

which for vanishing selection coefficients produces eq. (20).

#### A.5 Constant class proportions at birth

Until this point, we have defined fecundity rates as class-oriented quantities, and the rate at which an individual of class $j$ produces offspring of class $i$ was a variable portion of its total fecundity. In this section, we show that the analysis can be simplified under the additional assumption that one can write

$$f_{ij}(\boldsymbol{z}_j, \boldsymbol{x}_j(a), \boldsymbol{v}) = c_{ij} f_j(\boldsymbol{z}_j, \boldsymbol{x}_j(a), \boldsymbol{v}), \tag{A-65}$$

where $c_{ij}$ is the constant proportion of the offspring of individuals of class $j$ that are of class $i$ (i.e. such that $\sum_{i=1}^{n_c} c_{ij} = 1$).

##### A.5.1 Class-specific Hamiltonians and costates

Plugging eq. (A-65) into eq. (I-B), we have

$$\tilde{\boldsymbol{\lambda}}_{ij}(a) = c_{ij} \left. \frac{\partial \hat{R}_j(\boldsymbol{z}_j, \boldsymbol{x}_j(a), \boldsymbol{v}, a)}{\partial \boldsymbol{x}_j(a)} \right|_{\boldsymbol{z}=\boldsymbol{v}}, \tag{A-66}$$

where

$$\hat{R}_j(\boldsymbol{z}_j, \boldsymbol{x}_j(a), \boldsymbol{v}, a) = \int_a^\infty l_j(\tau) f_j(\boldsymbol{z}_j, \boldsymbol{x}_j(\tau), \boldsymbol{v}) \mathrm{d}\tau \tag{A-67}$$

is the expected remaining number of offspring of all classes for a mutant individual of class $j$. Let us now define the costate specific to parent class $j$ as the sums of class-oriented costates over offspring classes:

$$\tilde{\boldsymbol{\lambda}}_j(a) = \sum_{i=1}^{n_c} \tilde{\boldsymbol{\lambda}}_{ij}(a) = \left. \frac{\partial \hat{R}_j(\boldsymbol{z}_j, \boldsymbol{x}_j(a), \boldsymbol{v}, a)}{\partial \boldsymbol{x}_j(a)} \right|_{\boldsymbol{z}=\boldsymbol{v}}, \tag{A-68}$$

where the second line follows from using eq. (A-66). Likewise, let the Hamiltonian specific to parent class $j$ be the sum of class-oriented Hamiltonians over offspring classes:

$$\begin{aligned} H_j(\boldsymbol{z}_j, \boldsymbol{x}_j(a), \boldsymbol{v}) &= \sum_{i=1}^{n_c} H_{ij}(\boldsymbol{z}_j, \boldsymbol{x}_j(a), \boldsymbol{v}) \\ &= l_j(a) f_j(\boldsymbol{z}_j, \boldsymbol{x}_j(a), \boldsymbol{v}) \left( \sum_{i=1}^{n_c} c_{ij} \right) + \left( \sum_{i=1}^{n_c} \tilde{\boldsymbol{\lambda}}_{ij}(a) \right) \cdot \tilde{\boldsymbol{g}}_j(\boldsymbol{z}_j, \boldsymbol{x}_j(a), \boldsymbol{v}) \\ &= l_j(a) f_j(\boldsymbol{z}_j, \boldsymbol{x}_j(a), \boldsymbol{v}) + \tilde{\boldsymbol{\lambda}}_j(a) \cdot \tilde{\boldsymbol{g}}_j(\boldsymbol{z}_j, \boldsymbol{x}_j(a), \boldsymbol{v}), \end{aligned} \tag{A-69}$$

where the second and third lines follow from applying eq. (A-12) and eq. (A-65). Using eq. (A-68), and eq. (A-69), together with eq. (11), we can deduce that the components of $\tilde{\boldsymbol{\lambda}}_j(a)$ associated to survivorship and internal states, that is $\lambda_j^{\mathrm{l}}(a)$ and $\boldsymbol{\lambda}_j^{\mathrm{x}}(a)$ respectively, can be obtained as the solutions to the system of ODEs,

$$\begin{aligned}\frac{\mathrm{d}\lambda^{\mathrm{l}}_j(a)}{\mathrm{d}a} &= \sum_{i=1}^{n_\mathrm{c}} \frac{\mathrm{d}\lambda^{\mathrm{l}}_{ij}(a)}{\mathrm{d}a} = -\left.\frac{\partial H_j(\boldsymbol{z}_j, l_j(a), \boldsymbol{x}^\circ_j(a), \boldsymbol{v})}{\partial l_j(a)}\right|_{\boldsymbol{z}=\boldsymbol{v}} && \text{with i.c.} \quad \lambda^{\mathrm{l}}_j(0) = \sum_{i=1}^{n_\mathrm{c}} R^\circ_{ij}, \\ \frac{\mathrm{d}\boldsymbol{\lambda}^{\mathrm{x}}_j(a)}{\mathrm{d}a} &= \sum_{i=1}^{n_\mathrm{c}} \frac{\mathrm{d}\boldsymbol{\lambda}^{\mathrm{x}}_{ij}(a)}{\mathrm{d}a} = -\left.\frac{\partial H_j(\boldsymbol{z}_j, l^\circ_j(a), \boldsymbol{x}_j(a), \boldsymbol{v})}{\partial \boldsymbol{x}_j(a)}\right|_{\boldsymbol{z}=\boldsymbol{v}} && \text{with f.c.} \quad \lim_{a\to\infty} \boldsymbol{\lambda}^{\mathrm{x}}_j(a) = \mathbf{0}.\end{aligned} \tag{A-70}$$

In addition, from eqs. (A-66) and (A-68), it is clear that

$$\tilde{\boldsymbol{\lambda}}_{ij}(a) = c_{ij}\tilde{\boldsymbol{\lambda}}_j(a), \tag{A-71}$$

and thus that

$$H_{ij}(\boldsymbol{z}_j, \boldsymbol{x}_j(a), \boldsymbol{v}) = c_{ij} H_j(\boldsymbol{z}_j, \boldsymbol{x}_j(a), \boldsymbol{v}). \tag{A-72}$$

This means that in the case of constant class proportion at birth, the problem of solving the costates dynamics is reduced to solving $n_\mathrm{c}$ systems described by eq. (A-70). If the class-oriented components of selection, (e.g. $s_{j,l}(\boldsymbol{v})$) are still of biological interest, they can then be recovered using eqs. (A-71) and (A-72).

##### A.5.2 Fisher's condition and sexual systems

We now focus on a special case of constant class proportions delineated by two additional assumptions. First, we assume that the proportion of individuals of class $i$ within newborns is always constant (i.e. independent of the parent's identity); namely $c_{ij} = c_i$ for all $j$. In this case, element $(i, j)$ of the mutant next-generation matrix reads

$$\gamma_{ij} R_{ij}(\boldsymbol{z}_j, \boldsymbol{v}) = c_i \gamma_{ij} R_j(\boldsymbol{z}_j, \boldsymbol{v}), \tag{A-73}$$

where $R_j(\boldsymbol{z}_j, \boldsymbol{v}) = \hat{R}_j(\boldsymbol{z}_j, \boldsymbol{x}_j(0), \boldsymbol{v}, 0)$ (see eq. A-67) is the expected lifetime number of offspring of all classes produced by a mutant individual of class $j$. Second, we assume that each resident individual of class $j$ has, on average, one descendant individual of class $j$ in its lifetime, such that

$$R^\circ_{jj} = c_j R^\circ_j = 1. \tag{A-74}$$

While this condition has no intuitive biological interpretation in the general case, in obligate sexual populations with two classes (i.e. two sexes) it corresponds to 'Fisher's condition' (i.e. individuals must have a father and a mother), a well-known requirement for consistent sexual systems (Houston & McNamara 2006; Fromhage & Jennions 2016). We derive the following results for the general case of $n_\mathrm{c}$ classes.

Eq. (A-74) implies that $R^\circ_j = \frac{1}{c_j}$. Plugging this value into eq. (A-73) further implies that $\gamma_{ij} R^\circ_{ij} = c_i \gamma_{ij} \dfrac{1}{c_j}$. Making use of this property and defining the constant vector of class proportions $\mathbf{c} = (c_1, ..., c_j)$, the resident reproductive matrix can be written

$$\mathbf{R}^{\circ} = \mathbf{C}\Gamma\mathbf{C}^{-1}, \tag{A-75}$$

where $\mathbf{C} = \text{diag}(\mathbf{c})$ is the diagonal matrix of class frequencies a birth, $\mathbf{C}^{-1} = \text{diag}(1/\mathbf{c})$ is the diagonal matrix of inverse class frequencies, and where $\Gamma = [\gamma_{ij}]$ is the matrix of expected allelic transmission. Eq. (A-75) shows that $\mathbf{R}^{\circ}$ and $\Gamma$ are *similar* matrices (Caswell 2000; p.666), which means that they represent the same linear map in two different bases. This also means that the leading eigenvalue of $\Gamma$ must be the same as that of $\mathbf{R}^{\circ}$ (i.e. one) in any system matching this section's assumptions. For instance, this constraint rules out the case of haploid clonality, where $\Gamma$ is a matrix of ones with leading eigenvalue $n_c$.

It is possible to use the similarity between $\mathbf{R}^{\circ}$ and $\Gamma$ to facilitate the computation of $\boldsymbol{\nu}^{\circ}$ and $\boldsymbol{q}^{\circ}$. By definition of the eigenvectors and using the fact that $\mathbf{C}^{-1}\mathbf{C} = \mathbf{C}\mathbf{C}^{-1} = \mathbf{I}$, eq. (A-75) implies

$$\begin{cases} \boldsymbol{q}^{\circ} & = \mathbf{C}\Gamma\mathbf{C}^{-1}\boldsymbol{q}^{\circ} \\ \boldsymbol{\nu}^{\circ} & = \boldsymbol{\nu}^{\circ}\mathbf{C}\Gamma\mathbf{C}^{-1} \end{cases} \Rightarrow \begin{cases} (\mathbf{C}^{-1}\boldsymbol{q}^{\circ}) & = \Gamma(\mathbf{C}^{-1}\boldsymbol{q}^{\circ}) \\ (\boldsymbol{\nu}^{\circ}\mathbf{C}) & = (\boldsymbol{\nu}^{\circ}\mathbf{C})\Gamma \end{cases} \Rightarrow \begin{cases} \tilde{\boldsymbol{q}}^{\circ} & = \Gamma\tilde{\boldsymbol{q}}^{\circ} \\ \tilde{\boldsymbol{\nu}}^{\circ} & = \tilde{\boldsymbol{\nu}}^{\circ}\Gamma \end{cases}, \tag{A-76}$$

such that there is a simple relationship between $\boldsymbol{\nu}^{\circ}$ and $\boldsymbol{q}^{\circ}$ and the eigenvectors of $\Gamma$, $\tilde{\boldsymbol{\nu}}^{\circ}$ and $\tilde{\boldsymbol{q}}^{\circ}$,

$$\begin{aligned} \boldsymbol{q}^{\circ} &= \mathbf{C}\tilde{\boldsymbol{q}}^{\circ} \\ \boldsymbol{\nu}^{\circ} &= \tilde{\boldsymbol{\nu}}^{\circ}\mathbf{C}^{-1}. \end{aligned} \tag{A-77}$$

Eq. (A-77) is useful when $\Gamma$ has trivial eigenvectors under a given reproductive system. For instance, in a standard two-sex diploid system with a proportion $c$ and $(1-c)$ of males and females in newborns, all entries of $\Gamma$ are one half (i.e. each allele in both sexes has half a chance to be transmitted to a given offspring), such that it admits eigenvectors $\tilde{\boldsymbol{q}}^{\circ} = (1,1)$ and $\tilde{\boldsymbol{\nu}}^{\circ} = (1,1)$. Using eq. (A-77) and scaling gives

$$\begin{aligned} \boldsymbol{q}^{\circ} &= (c, 1-c) \\ \boldsymbol{\nu}^{\circ} &= (\frac{1}{2c}, \frac{1}{2(1-c)}). \end{aligned} \tag{A-78}$$

Under haplodiploidy, we have

$$\Gamma = \begin{pmatrix} 0 & 1/2 \\ 1 & 1/2 \end{pmatrix} \tag{A-79}$$

which admits eigenvectors $\tilde{\boldsymbol{q}}^{\circ} = (1/2, 1)$ and $\tilde{\boldsymbol{\nu}}^{\circ} = (1,1)$. Using eq. (A-77) and scaling gives

$$\begin{aligned} \boldsymbol{q}^{\circ} &= (\frac{c}{2-c}, \frac{2(1-c)}{2-c}) \\ \boldsymbol{\nu}^{\circ} &= (\frac{2-c}{3c}, \frac{2-c}{3(1-c)}). \end{aligned} \tag{A-80}$$

Eqs. (A-78) and (A-80) illustrate the correspondence of $\boldsymbol{q}^{\circ}$ with allelic class frequencies at birth, including asymmetries due to asymmetric allelic transmission and ploidy.

#### A.6 Analysis of selection on genetically correlated traits

In this section we describe how to adapt our results to study cases where some traits are genetically correlated, i.e. share a unique genetic value and cannot be affected separately by mutation. For ease of presentation, we describe only the special case where a single pair of traits displays such genetic constraints, while other traits evolve independently. The rationale in this section, however, can readily be extended to several pairs, or to groups, of genetically correlated traits. In the main text, we use the approach described here to study the case of correlated growth rates as an extension to our example model.

Assume two genetically correlated traits $(i, k)$ and $(j, l)$, expressed in different classes ($i \neq j$) or in the same class ($i = j$). Since due to genetic constraints we have $\eta_{i,k} = \eta_{j,l}$ for all mutations, this is a akin to having a single evolving trait expressed in two different contexts. In that case, mutant and resident trait values for correlated traits must be equal ($u_{i,k} = u_{j,l}$ and $v_{i,k} = v_{j,l}$) and eq. (21) must be satisfied for the first-order uninvadibility condition described by eq. (4) to be met at $\boldsymbol{v}^*$, under the constraint that $v^*_{i,k} = v^*_{j,l}$. In turn, evaluating the convergence stability and uninvadibility of such a singular trait requires a new Jacobian matrix $\check{\mathbf{J}}(\boldsymbol{v}^*)$ and a new Hessian matrix $\check{\mathbf{H}}(\boldsymbol{v}^*)$, both with one less row and column (i.e. one less independently evolving trait) than the matrices $\mathbf{J}(\boldsymbol{v}^*)$ and $\mathbf{H}(\boldsymbol{v}^*)$ described in the main text (eq. 6).

The matrix $\check{\mathbf{J}}(\boldsymbol{v}^*)$ can be obtained by first computing $\mathbf{J}(\boldsymbol{v}^*)$, where $\boldsymbol{v}^*$ is the singular trait satisfying eq. (21), before replacing its pair of rows $(i, k)$ and $(j, l)$ by a single row $(j, l)$ that is their element-wise sum, and its pair of columns $(i, k)$ and $(j, l)$ by a single column $(j, l)$ that is their element-wise sum. This operation brings together elements of $\mathbf{J}(\boldsymbol{v}^*)$ that pertain to the effect of each resident traits on both correlated traits (i.e. rows $(i, k)$ and $(j, l)$), and elements of $\mathbf{J}(\boldsymbol{v}^*)$ that pertain to the effect of both correlated resident traits on each directional selection coefficient (i.e. columns $(i, k)$ and $(j, l)$). More formally, the element $(m, n)$ of the newly created row $(j, l)$ of $\check{\mathbf{J}}(\boldsymbol{v}^*)$ is

$$\check{j}_{jm,ln} = \left( \frac{\partial s_{i,k}(\boldsymbol{v})}{\partial v_{m,n}} + \frac{\partial s_{j,l}(\boldsymbol{v})}{\partial v_{m,n}} \right) \Bigg|_{\boldsymbol{v}=\boldsymbol{v}^*}, \tag{A-81}$$

and the element $(m, n)$ of the newly created column $(j, l)$ of $\check{\mathbf{J}}(\boldsymbol{v}^*)$ is

$$\check{j}_{mj,nl} = \left( \frac{\partial s_{m,n}(\boldsymbol{v})}{\partial v_{i,k}} + \frac{\partial s_{m,n}(\boldsymbol{v})}{\partial v_{j,l}} \right) \Bigg|_{\boldsymbol{v}=\boldsymbol{v}^*}, \tag{A-82}$$

for all traits $(m, n) \neq (i, k), (j, l)$. The newly created diagonal element $(jj, ll)$ of $\check{\mathbf{J}}(\boldsymbol{v}^*)$, in turn, contains the effects of both correlated resident traits on both $s_{i,k}(\boldsymbol{v})$ and $s_{j,l}(\boldsymbol{v})$, and reads

$$\check{\jmath}_{jj,ll} = \left( \frac{\partial s_{i,k}(\boldsymbol{v})}{\partial v_{i,k}} + \frac{\partial s_{i,k}(\boldsymbol{v})}{\partial v_{j,l}} + \frac{\partial s_{j,l}(\boldsymbol{v})}{\partial v_{i,k}} + \frac{\partial s_{j,l}(\boldsymbol{v})}{\partial v_{j,l}} \right)\Bigg|_{\boldsymbol{v}=\boldsymbol{v}^*}. \tag{A-83}$$

Similarly, the matrix $\check{\mathbf{H}}(\boldsymbol{v}^*)$ can be obtained by first computing $\mathbf{H}(\boldsymbol{v}^*)$, where $\boldsymbol{v}^*$ is the singular trait satisfying eq. (21), before replacing its pair of rows $(i,k)$ and $(j,l)$ by a single row $(j,l)$ that is their element-wise sum, and its pair of columns $(i,k)$ and $(j,l)$ by a single column $(j,l)$ that is their element-wise sum. This operation brings together elements of $\mathbf{H}(\boldsymbol{v}^*)$ that pertain to the multiplicative effect of both correlated traits with the same third trait. More formally, the element $(m,n)$ of the newly created row $(j,l)$ of $\check{\mathbf{H}}(\boldsymbol{v}^*)$ is

$$\check{h}_{jm,ln}(\boldsymbol{v}^*) = h_{im,kn}(\boldsymbol{v}^*) + h_{jm,ln}(\boldsymbol{v}^*), \tag{A-84}$$

and the element $(m,n)$ of the newly created column $(j,l)$ of $\check{\mathbf{H}}(\boldsymbol{v}^*)$ is

$$\check{h}_{mj,nl}(\boldsymbol{v}^*) = h_{mi,nk}(\boldsymbol{v}^*) + h_{mj,nl}(\boldsymbol{v}^*), \tag{A-85}$$

for all traits $(m,n) \neq (i,k),(j,l)$. The newly created diagonal element $(jj,ll)$ of $\check{\mathbf{H}}(\boldsymbol{v}^*)$, in turn, collects the multiplicative effects of both traits together and the multiplicative effects of both trait independently, that is

$$\check{h}_{jj,ll}(\boldsymbol{v}^*) = h_{ij,kl}(\boldsymbol{v}^*) + h_{ji,lk}(\boldsymbol{v}^*) + h_{jj,ll}(\boldsymbol{v}^*) + h_{ii,kk}(\boldsymbol{v}^*). \tag{A-86}$$

The sums in eqs. (A-84)–(A-86) can be computed separately for each component of selection described in eq. (15) to recover these components under genetic constraints. Note that when computing these equations, the term described in eq. (A-30) must be included. This is because in the case of genetically correlated traits we have $s_{i,k}(\boldsymbol{v}^*) \neq 0$ and $s_{j,l}(\boldsymbol{v}^*) \neq 0$, according to eq. (21). For the same reason, the perturbations of allelic frequencies in eq. (16f) (i.e. $\partial \boldsymbol{q}(\boldsymbol{z},\boldsymbol{v}^*)/\partial z_{j,l}$) should be computed using eq. (A-64) instead of eq. (20).

### Appendix B - Example model

#### B.1 Resident quantities

In this section, we detail the derivations of the resident quantities used in the example model of section 4.1. We start with the fact that at equilibrium, that is when the number $n_j^{\circ}(a)$ of individuals of sex $j$ and age $a$ is constant, the population structure satisfies the equation

$$n_j^{\circ}(a) = c_j l_j^{\circ}(a) \int_0^{\infty} n_{\mathrm{f}}^{\circ}(s) f_{\mathrm{f}}(x_{\mathrm{f}}^{\circ}(s), \boldsymbol{v}) \mathrm{d}s, \tag{B-87}$$

where $c_{\rm f} = (1-c)$ and $c_{\rm m} = c$. Eq. (B-87) means that each individual of age $a$ and sex $j$ must have been born to a female of any age $s$ (i.e. with class-oriented fecundity $c_j f_{\rm f}(x_{\rm f}^{\circ}(s), \boldsymbol{v})$) and must then have survived to age $a$ (the integral in eq. (B-87) is the total number of newborns produced in the population).

##### B.1.1 Age distribution

From eq. (B-87), the probability density $\tilde{\iota}_j^{\circ}(a)$ of individuals of age $a$, *given* that they are of sex $j$ (i.e. such that $\int_0^{\infty} \tilde{\iota}_j^{\circ}(a) \mathrm{d}a = 1$), can be obtained as

$$\tilde{\iota}_j^{\circ}(a) = \frac{n_j^{\circ}(a)}{\int_0^{\infty} n_j^{\circ}(\tau) \mathrm{d}\tau} = \frac{l_j^{\circ}(a)}{\int_0^{\infty} l_j^{\circ}(\tau) \mathrm{d}\tau} = \mu_j^{\circ} l_j^{\circ}(a), \tag{B-88}$$

where eq. (25) was used. Eq. (B-88) is biologically consistent as it implies that $\tilde{\iota}_j^{\circ}(0) = \mu_j^{\circ}$, meaning that the probability density of newborns matches the death rate within sex $j$. Also from eq. (B-87), the probability density $\iota_j^{\circ}(a)$ of individuals of sex $j$ and age $a$ in the total resident population (i.e. such that $\int_0^{\infty} \iota_{\rm m}^{\circ}(a) + \iota_{\rm f}^{\circ}(a) \mathrm{d}a = 1$) can be obtained as

$$\iota_j^{\circ}(a) = \frac{n_j^{\circ}(a)}{\int_0^{\infty} n_{\rm f}^{\circ}(\tau) + n_{\rm m}^{\circ}(\tau) \mathrm{d}\tau} = \frac{c_j l_j^{\circ}(a)}{\int_0^{\infty} (1-c) l_{\rm f}^{\circ}(a) + c l_{\rm m}^{\circ}(a) \mathrm{d}\tau} = \frac{\mu_{\rm f}^{\circ} \mu_{\rm m}^{\circ}}{(1-c)\mu_{\rm m}^{\circ} + c\mu_{\rm f}^{\circ}} c_j l_j^{\circ}(a). \tag{B-89}$$

From eq. (B-89), the secondary sex-ratio (given in eq. 30) can be obtained as

$$\tilde{c}^{\circ} = \int_0^{\infty} \iota_{\rm m}^{\circ}(a) \mathrm{d}a = \frac{c\mu_{\rm f}^{\circ} \mu_{\rm m}^{\circ}}{(1-c)\mu_{\rm m}^{\circ} + c\mu_{\rm f}^{\circ}} \int_0^{\infty} l_{\rm m}^{\circ}(a) \mathrm{d}a = \frac{c\mu_{\rm f}^{\circ}}{(1-c)\mu_{\rm m}^{\circ} + c\mu_{\rm f}^{\circ}}. \tag{B-90}$$

##### B.1.2 Population density

Dividing eq. (B-87) by $\int_0^{\infty} n_{\rm f}^{\circ}(\tau) + n_{\rm m}^{\circ}(\tau) \mathrm{d}\tau$ and using eq. (B-89), implies that

$$\iota_{\rm f}^{\circ}(0) = (1-c) \int_0^{\infty} \iota_{\rm f}^{\circ}(a) f_{\rm f}(x_{\rm f}^{\circ}(a), \boldsymbol{v}) \mathrm{d}a. \tag{B-91}$$

Substituting eq. (B-89) and eq. (27) into eq. (B-91) and simplifying, we obtain

$$1 = \left(1 - \frac{N_{\rm T}^{\circ}}{K}\right)(1-c) \int_0^{\infty} l_{\rm f}^{\circ}(a) x_{\rm f}^{\circ}(a) \mathrm{d}a. \tag{B-92}$$

Eq. (B-92) is demographically consistent since it entails that at equilibrium, each resident female produces an average of one new female during its lifetime (i.e. such that their total number is fixed). From eq. (B-92), we have

$$N_{\rm T}^{\circ} = K \left[1 - \frac{1}{(1-c)\int_0^{\infty} l_{\rm f}^{\circ}(a) x_{\rm f}^{\circ}(a) \mathrm{d}a}\right]. \tag{B-93}$$

##### B.1.3 Size distribution

Here we derive the expression for the frequency $\chi_j^\circ(x)$ of individuals of size $x$ within sex $j$ in the resident population at equilibrium (eq. (26)). To that aim, we note that eq. (13) from De Roos (1997) provides a partial differential equation describing the time dynamics of the density function of any internal state satisfying our main text eq. (12) in a monomorphic population. With our notation, and denoting $\chi_j^\circ(x,t)$ the probability density of individuals of size $x$ within sex $j$ at an arbitrary time $t$, eq. (13) from De Roos (1997) entails that

$$\begin{aligned}\frac{\partial \chi_j^\circ(x,t)}{\partial t} &= -\frac{\partial[(v_j - \alpha x)\chi_j^\circ(x,t)]}{\partial x} - \mu_j(v_j, x, \boldsymbol{v})\chi_j^\circ(x,t) \\ &= \alpha\chi_j^\circ(x,t) - (v_j - \alpha x)\frac{\partial \chi_j^\circ(x,t)}{\partial x} - \mu_j(v_j, x, \boldsymbol{v})\chi_j^\circ(x,t).\end{aligned} \tag{B-94}$$

At demographic equilibrium, the distribution of size in sex $j$ is at a steady state, such that $\partial\chi_j^\circ(x,t)/\partial t = 0$ and $\chi_j^\circ(x,t) = \chi_j^\circ(x)$ for all $t$. Simplifying eq. (B-94) accordingly, we obtain

$$\frac{\partial \chi_j^\circ(x)}{\partial x} = \frac{\alpha - \mu_j^\circ}{v_j - \alpha x}\chi_j^\circ(x), \tag{B-95}$$

which is an ODE with solution

$$\chi_j^\circ(x) = \chi_j^\circ(0)\left(\frac{v_j}{v_j - \alpha x}\right)^{\frac{\alpha - \mu_j^\circ}{\alpha}}. \tag{B-96}$$

By integrating eq. (B-96) over the range of possible sizes (i.e. between 0 and $v_j/\alpha$, according to eq. 25), we find that

$$\int_0^\infty \chi_j^\circ(x)\mathrm{d}x = \frac{v_j\chi_j^\circ(0)}{\mu_j^\circ}. \tag{B-97}$$

For this expression to be equal to 1 (i.e. as required from a probability density function), it must be that $\chi_j^\circ(0) = \mu_j^\circ/v_j$. Plugging this value into eq. (B-96), we obtain

$$\chi_j^\circ(x) = \frac{\mu_j^\circ}{v_j}\left(\frac{v_j}{v_j - \alpha x}\right)^{\frac{\alpha - \mu_j^\circ}{\alpha}}. \tag{B-98}$$

Using eq. (B-98), we can compute the expectation (i.e. average) and the variance of the size distribution in sex $j$ at equilibrium, as

$$\overline{x_j^\circ} = \int_0^\infty \iota_j^\circ(a)x_j^\circ(a)\mathrm{d}a = \int_0^{\tilde{x}_j^\circ} \chi_j^\circ(x)x\mathrm{d}x = \frac{v_j}{\alpha + \mu_j^\circ}, \tag{B-99}$$

and

$$\int_0^\infty \iota_j^\circ(a)(x_j^\circ(a) - \overline{x_j^\circ})^2\mathrm{d}a = \int_0^{\tilde{x}_j^\circ} \chi_j^\circ(x)(x - \overline{x_j^\circ})^2\mathrm{d}x = \frac{v_j^2\mu_j^\circ}{(\alpha + \mu_j^\circ)^2(2\alpha + \mu_j^\circ)}, \tag{B-100}$$

respectively.

#### B.2 ESS in the absence of class-structure and ontogeny

Consider a simple population with a single haploid class, where individuals have a single constant (i.e. static) size $x$. Assume that size is directly determined by a single trait $z$ (i.e. $x = z$), and determines absolute fecundity (i.e. $f(x) = x$). Under the same assumptions and approach as in the main text, the basic reproductive number of a mutant with size $x = z = u$ in this model is

$$\begin{aligned} R(u, v) &= \int_0^\infty l(a) u \left(1 - \frac{N_\mathrm{T}^\circ}{K}\right) \mathrm{d}a \\ &= \frac{u}{\mu(u)} \frac{K - N_\mathrm{T}^\circ}{K}, \end{aligned} \tag{B-101}$$

where $l(a) = e^{-\mu(u)a}$, $\mu(u) = \mu_e + \beta u^2$ and where

$$N_\mathrm{T}^\circ = K \left(1 - \frac{\mu^\circ}{v}\right) \tag{B-102}$$

is the equilibrium resident population density, obtained as the solution to $R(v, v) = 1$. With this, the directional selection pressure acting on trait $z$ is

$$s(v) = \left.\frac{\partial R(u, v)}{\partial u}\right|_{u=v} = \frac{1}{v} \left(\frac{\mu_\mathrm{e} - \beta v^2}{\mu_\mathrm{e} + \beta v^2}\right). \tag{B-103}$$

This entails a single singular trait value

$$v^* = \sqrt{\frac{\mu_\mathrm{e}}{\beta}}, \tag{B-104}$$

which balances growth and survival. This result can be directly recovered from equation eq. (36). On the one hand, according to eq. (I-B) and using eq. (B-93) with $\alpha = 0$, in the case of a static state and a single class we have

$$\lambda^\mathrm{x}(a) = \left.\frac{\partial \int_a^\infty l(\tau) x (1 - \frac{N_\mathrm{T}^\circ}{K}) \mathrm{d}\tau}{\partial x}\right|_{\boldsymbol{z}=\boldsymbol{v}} = \int_a^\infty l^\circ(\tau) \left(1 - \frac{N_\mathrm{T}^\circ}{K}\right) \mathrm{d}\tau = \frac{l^\circ(a)\mu^\circ}{v}, \tag{B-105}$$

such that

$$\int_0^\infty \lambda^\mathrm{x}(a) \mathrm{d}a = \frac{1}{v}. \tag{B-106}$$

On the other hand, according to eq. (I-C), we have

$$\lambda^\mathrm{l}(a) l^\circ(a) = \int_a^\infty l^\circ(\tau) v \left(1 - \frac{N_\mathrm{T}^\circ}{K}\right) \mathrm{d}\tau = l^\circ(a), \tag{B-107}$$

which shows that at equilibrium and when fecundity is constant, the expected remaining reproductive output corresponds to survivorship. According to eq. (B-107), we have

$$\int_0^\infty \lambda^\mathrm{l}(a) l^\circ(a) \mathrm{d}a = \frac{1}{\mu^\circ}. \tag{B-108}$$

Plugging eq. (B-106) and eq. (B-107) into eq. (36), we obtain the equation

$$v^* = \frac{1}{2\beta}\frac{\mu_{\mathrm{e}} + \beta {v^*}^2}{v^*}, \tag{B-109}$$

of which eq. (B-104) is the only positive solution.

It is straightforward to show that the singular trait in eq. (B-104) is both convergence stable and uninvadable. From eq. (B-103) we have

$$\left.\frac{\partial s(v)}{\partial v}\right|_{v=v^*} = -\frac{\beta}{\mu_e}, \tag{B-110}$$

showing the former, while from eq. (B-101) we have

$$\left.\frac{\partial^2 R(u,v)}{\partial u^2}\right|_{u=v=v^*} = -\frac{\beta}{\mu_e}, \tag{B-111}$$

showing the latter.

Thus, in the absence of sexes and ontogeny, our model reduces to a simple trade-off between fecundity and survival, with the long-term stable investment into fecundity being higher under higher extrinsic mortality, and lower under higher survival costs. Because this case removes mating interactions and variance in resident states, no polymorphism can evolve.

#### B.3 Individual-based simulations

Here we describe the individual-based simulations used in fig. 4 (as well as fig. S3 for haplodiploids, and fig. S5 for genetically correlated traits). Python scripts implementing these simulations are available at zenodo.org (DOI: 10.5281/zenodo.17252309).

Our simulations rely on a discrete-time approximation of the continuous-time life-cycle described in the main text, with small time steps of fixed length $\tau$. Growth and mortality dynamics are approximated using Euler's method (Süli & Mayers 2003; p.341). The number of offspring produced over one time step by a female with fecundity $f$ at the start of the interval is drawn from a Poisson distribution with rate $f\tau$. This is equivalent to assuming that, during the considered interval, the waiting time to the next offspring is exponentially distributed with parameter $f$ (Last & Penrose 2017; theorem 7.2, p. 59). For a small time step $\tau$, the discrete-time model should thus approximate the continuous-time process.

Each individual in the simulation model is characterized by its age, its size, its sex and its genotype. Except for haploid males in the haplodiploid version, each individual is diploid and carries a pair of haplotypes. Each haplotype consists of two real values, giving the allelic values it encodes for male and female basal growth rates, respectively. Upon birth, the trait $z$ of each individual is calculated as the average of the allelic values it carries and that correspond to its sex. The sex of each individual is drawn at random, with a probability $c$ of being male.

Starting with a population where all individuals have age and size 0, and with haplotypes all set to the same arbitrary pair of values (i.e. we start with a monomorphic population and set the initial population size to 4000), the population is then iterated through the following steps:

1. **Aging and growing**. Each individual sets its age $a$ to $a + \tau$, and its size $x$ to $x + (z - \alpha x)\tau$ (according to eq. 22).
2. **Dying**. Each individual of sex $j$ dies with probability $(\mu_{\mathrm{e},j} + \beta z^2)\tau$ (according to eq. 23). Dead individuals are removed from the population.
3. **Mating**. For each female of size $x_{\mathrm{f}}$ in the population, a sample is drawn from a Poisson distribution with rate $x_{\mathrm{f}}(1 - N_{\mathrm{T}}/K)\tau$ (see eq. 27), which represents the number of offspring that this female produced in the current time step. For each of these offspring, a father is sampled (with replacement) from the whole population of males. There, the probability for a male of size $x_{\mathrm{m}}$ to be drawn is proportional to $e^{-\kappa(x_{\mathrm{m}}-x_{\mathrm{f}})^2}$ (i.e. its absolute competitive ability, see main text).
4. **Recombination and mutation**. For each offspring in the population, a genotype is constructed from that of its parents. Each parent produces a gamete, which is a free recombinant of its own two haplotypes (i.e. probability to recombine is 0.5). Once gametes are produced, each allelic value on each gamete can mutate with probability 0.01. Mutation effects are drawn in a normal distribution with mean 0 and standard deviation 0.1.
5. **Fertilization**. Each offspring is instantiated with age 0 and size 0, and with its two parental gametes as its genotype. Sex is drawn at random, with a probability $c$ of being male. In the special case of newborn males in haplodiploid simulations, both haplotypes are set as the mother's gamete (a valid trick under additive genetic effects).
